# A high-resolution model of gene expression during *Gossypium hirsutum* (cotton) fiber development

**DOI:** 10.1101/2024.07.20.604417

**Authors:** Corrinne E Grover, Josef J Jareczek, Sivakumar Swaminathan, Youngwoo Lee, Alexander H Howell, Heena Rani, Mark A Arick, Alexis G Leach, Emma R Miller, Pengcheng Yang, Guanjing Hu, Xianpeng Xiong, Eileen L Mallery, Daniel G Peterson, Jun Xie, Candace H Haigler, Olga A Zabotina, Daniel B Szymanski, Jonathan F Wendel

## Abstract

Cotton fiber development relies on complex and intricate biological processes to transform newly differentiated fiber initials into the mature, extravagantly elongated cellulosic cells that are the foundation of this economically important cash crop. Here we extend previous research into cotton fiber development by employing controlled conditions to minimize variability and utilizing time-series sampling and analyses to capture daily transcriptomic changes from early elongation through the early stages of secondary wall synthesis (6 to 24 days post anthesis; DPA). A majority of genes are expressed in fiber, largely partitioned into two major coexpression modules that represent genes whose expression generally increases or decreases during development. Differential gene expression reveals a massive transcriptomic shift between 16 and 17 DPA, corresponding to the onset of the transition phase that leads to secondary wall synthesis. Subtle gene expression changes are captured by the daily sampling, which are discussed in the context of fiber development. Coexpression and gene regulatory networks are constructed and associated with phenotypic aspects of fiber development, including turgor and cellulose production. Key genes are considered in the broader context of plant secondary wall synthesis, noting their known and putative roles in cotton fiber development. The analyses presented here highlight the importance of fine-scale temporal sampling on understanding developmental processes and offer insight into genes and regulatory networks that may be important in conferring the unique fiber phenotype.

## Introduction

Cotton fibers are individual cells that emerge from the developing ovule epidermis and develop over a period of about two months from initiation to maturity. Fiber development entails a tightly coordinated series of overlapping stages that oversee the transformation of individual cells from spherical epidermal protrusions on the ovular surface to mature fibers whose length can exceed 5 cm and whose cell wall (CW) composition approaches 98% cellulose (Kim and Triplett, 2001; Butterworth *et al*., 2009; Kim, 2018; Jareczek, Grover and Wendel, 2023). These highly polarized cells are both useful models for plant cell morphogenesis (Kim and Triplett, 2001; Butterworth *et al*., 2009; Kim, 2018; Jareczek, Grover and Wendel, 2023; Haigler *et al*., 2012) and form the foundation of a multibillion dollar textile industry; therefore, understanding their growth and development are important from both agronomic and fundamental biology perspectives. Although four species of cotton have been independently domesticated, *Gossypium hirsutum* (or Upland cotton), comprises the vast majority of the market share (∼95%) due to its high yield, greater pest resistance, and environmental adaptability (Constable *et al*., 2015). *Gossypium hirsutum* is an allopolyploid containing two coresident genomes (At, Dt) donated by the progenitor diploids at the time of polyploid formation circa 1 million years ago (reviewed in (Viot and Wendel, 2023; Hu, Grover, Yuan, *et al*., 2021)). Following its initial domestication, *G. hirsutum* experienced strong directional selection for intensely elongated fiber (Kim, 2015; Applequist *et al*., 2001), among other traits, which resulted in massive reorganization of the fiber transcriptome and tighter coordination among fiber-related genes (Gallagher *et al*., 2020; Rapp *et al*., 2010).

At the biosynthetic level, fiber development requires intricate coordination of cellular processes that establish the shape and length of the fiber cell. Morphogenesis takes place over several overlapping stages (Figure 1) whose interplay ultimately determines fiber characteristics. The first stage, initiation, begins on the ovular surface around the time of anthesis (i.e., flower opening) and is regulated by phytohormones (e.g., positive regulators include auxin, brassinosteroids, and jasmonic acid; reviewed in (Xiao *et al*., 2019; Jareczek, Grover and Wendel, 2023), as well as reactive oxygen species (ROS). Evolutionarily conserved MYB cell-fate control genes are implicated in fiber initiation, as are many other genes (L., Wang *et al*., 2021; Zou *et al*., 2022; Qin *et al*., 2019; N.,-N., Wang *et al*., 2021; Ando *et al*., 2021; Zhao *et al*., 2019; Jiao *et al*., 2023; Jiang *et al*., 2021; Hu *et al*., 2016), including those involved in cytoskeleton-dependent cell wall patterning (Li *et al*., 2005; Qu *et al*., 2012; Gilbert *et al*., 2014; Y., Zhang *et al*., 2021). Fiber cells elongate through a highly polarized form of anisotropic diffuse growth over about 3 weeks (Kim, 2018). Soon after fiber elongation begins, the cells taper under the influence of apical microtubules and cellulose to progressively reduce and restrict cell diameter throughout development (Stiff and Haigler, 2016; Yanagisawa *et al*., 2015; Yanagisawa *et al*., 2022). This specialized apical domain and transverse network of microtubules help to establish fiber cell diameter and enable resistance to swelling along the cell axis as elongation continues. Transverse cortical microtubules direct the synthesis of parallel stiff cellulose microfibrils that resist radial expansion as high turgor pressure drives anisotropic growth (Lockhart, 1965; Proseus *et al*., 2000; Ryser, 1977; Tiwari and Wilkins, 1995; Qin and Zhu, 2011; Yu *et al*., 2019).

**Figure 1.**
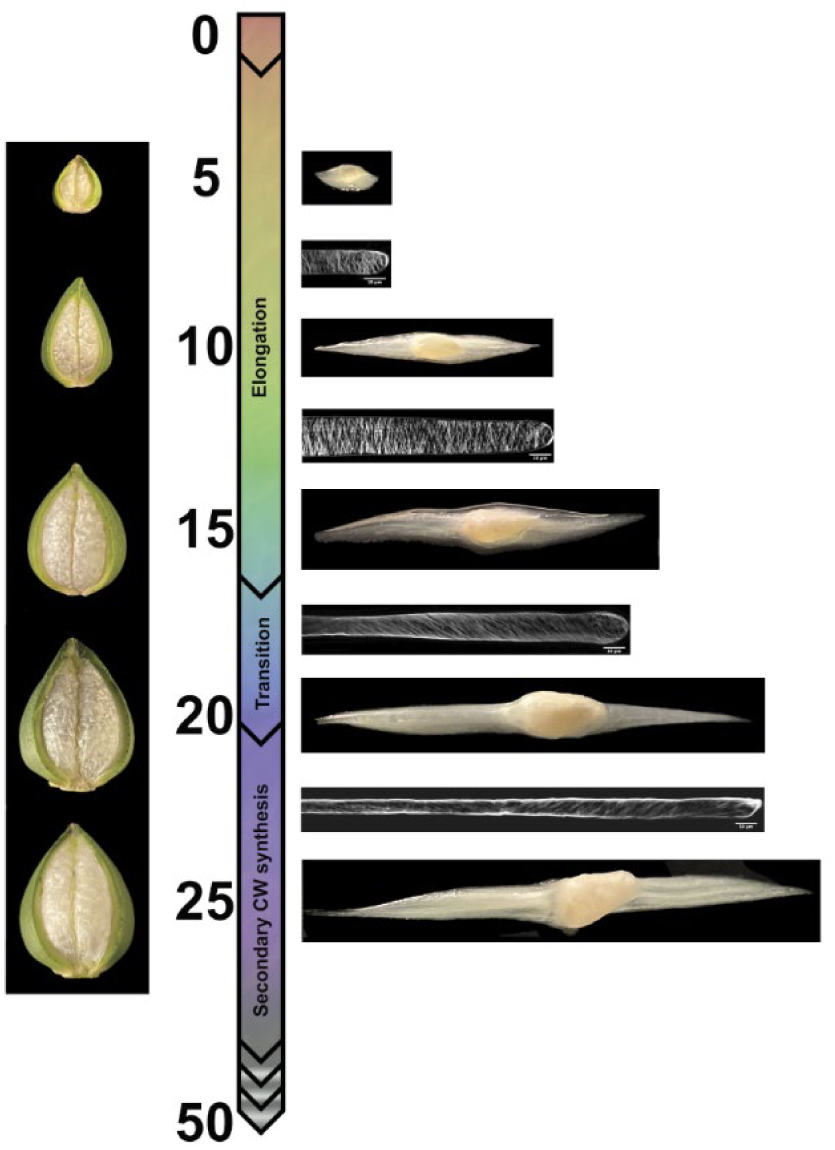
Illustration of cotton fiber developmental timeline focusing on the first half of development. Cotton fiber development starts with initiation of fiber cells on the ovule seed coat, which begins around the time of flower opening (anthesis) and continues during the first few days of seed development during which the fiber cells taper to reduce cell diameter (by 2 days post anthesis; DPA). The elongation phase, which includes primary wall synthesis, has complex dynamics and persists for about 20 days. At approximately 16 DPA, the transition between elongation and cell wall thickening begins. The mature, cellulose-rich fiber is fully formed at about 50 DPA. Images are placed in their approximate position along the developmental timeline. Cut capsules (“bolls”) and developing fibers are shown to the left and right of the timeline, respectively. Confocal images of developing fibers show the orientation of the cellulose microfibrils, which changes from approximately transverse during elongation to an increasingly steep helix beginning at the transition stage. Images of growing ovules with fiber combed away in two directions are intercalated.

The composition and material properties of the cell wall matrix polysaccharides are also tuned during the elongation phase to enable predictable cell shape outcomes (Avci *et al*., 2013; Swaminathan *et al*., 2024; Delmer *et al*., 2024). Complex interactions between the cellulose and matrix components of the wall likely underlie much of the observed growth rate variability (Yanagisawa *et al*., 2015; Yanagisawa *et al*., 2022). Important polysaccharides during this phase are those such as cellulose, xyloglucan, and pectin (Haigler *et al*., 2012; Kim and Triplett, 2001; Avci *et al*., 2013; Pettolino *et al*., 2022; Jareczek, Grover and Wendel, 2023), whose arrangement and composition results in unidirectional extensibility. Both turgor and cell wall stiffness influence fiber growth rate (Yanagisawa *et al*., 2015; Yanagisawa *et al*., 2022), which makes turgor modulation and fiber cell wall composition change during development (Avci *et al*., 2013; Meinert and Delmer, 1977; Pettolino *et al*., 2022) active areas of research.

As with the initiation phase, numerous genes have been implicated in elongation, including transcription factors and various cytoskeletal genes (Pu *et al*., 2008; Machado *et al*., 2009; Shan *et al*., 2014; Luo *et al*., 2007; Yang *et al*., 2014; Zhang *et al*., 2017; Huang *et al*., 2021; Takatsuka *et al*., 2018). Phytohormones continue to play an important role in elongation (Jareczek, Grover and Wendel, 2023), with many ethylene biosynthetic genes and pathways upregulated during this stage (Ahmed *et al*., 2018; Xiao *et al*., 2019). These in turn influence the expression of fiber-related genes such as cellulose synthase, expansins, and sucrose synthase, while also influencing both the brassinosteroid pathway and ROS management, the latter contributing to anisotropic growth in the fiber (Ahmed *et al*., 2018; Xiao *et al*., 2019; Tang *et al*., 2014; Jareczek, Grover and Wendel, 2023).

A major developmental transition takes place somewhere between ∼16 to 20 DPA (Figure 1), marking the switch from fiber elongation to secondary cell wall (SCW) synthesis (Meinert and Delmer, 1977). The transition is a distinct development stage characterized by: increased cellulose synthesis; changes in microtubule and cellulose microfibril orientation; decreased synthesis of primary cell wall (PCW) polysaccharides; and degradation of the cotton fiber middle lamella (CFML), among other changes in biochemical and cellular features (Haigler *et al*., 2012; Singh *et al*., 2009). Correspondingly extensive changes in gene expression and other regulatory processes (e.g., phytohormone activity) occur (Zhou *et al*., 2019; Tuttle *et al*., 2015; Jareczek, Grover and Wendel, 2023). The fiber, which is composed of 90 - 95% cellulose at maturity, commits increasing resources toward cellulose production as the fiber moves into the last phase of SCW thickening (∼23 DPA to 45 DPA; Figure 1). Some of the regulatory genes involved in the transition include NAC-domain factors (e.g., SND1 and TALE family genes; (Zhong *et al*., 2006; Ma *et al*., 2019)), MYB genes (including GhMYBL1; (Zhong *et al*., 2006; Li *et al*., 2009; Sun *et al*., 2015)), and the transcription factor Hot216, a KIP-related protein that regulates a network of ∼1000 cell wall synthesis genes (Li *et al*., 2020). As the cell moves into SCW synthesis, a subgroup of cellulose synthases become highly expressed (Kim, 2018), along with genes related to regulation of UDP-glucose, the substrate for synthesis of cellulose and some other cell wall polymers (Buchala, 1999). Many other genes are also up-regulated, given the complex changes in the metabolome during the SCW stage (Tuttle *et al*., 2015).

The molecular underpinnings of fiber development and various fiber properties (e.g., length, strength) in *G. hirsutum* have been evaluated at the transcriptome level using different comparative strategies and time points. Many comparisons have evaluated the expression differences that underlie important fiber morphologies via differential gene expression at key timepoints between accessions that vary in these important fiber properties (Qin *et al*., 2019; Li *et al*., 2023; Naoumkina *et al*., 2015; Islam *et al*., 2016) or among time points sampled (Wang *et al*., 2010; Gallagher *et al*., 2020; Jareczek, Grover, Hu, *et al*., 2023; Yoo and Wendel, 2014), resulting in many of the insights mentioned above. Others have made interspecific comparisons to *G. barbadense*, whose fiber possesses several desirable properties (Tuttle *et al*., 2015; Jareczek, Grover, Hu, *et al*., 2023; Zhu *et al*., 2011; Chen *et al*., 2012; Rapp *et al*., 2010), or compared developmental timelines between wild and domesticated forms of *G. hirsutum (Jareczek, Grover, Hu, et al., 2023; Rapp et al., 2010; Gallagher et al., 2020; Yoo and Wendel, 2014)*. The emerging picture from these and other studies is that fiber development is transcriptionally complex, in part reflecting overlap and compromises among the gene networks regulating important fiber properties such as length and strength.

In this study, we extend our prior understanding of fiber development by sampling the transcriptome more densely than in prior studies and, combined with data from other ‘omics’ and fiber phenotypes, provide preliminary information regarding the networks controlling cotton fiber development. These data allow a more fine-scale characterization of elongation, the transition phase, and SCW synthesis, when fiber becomes increasingly committed to cellulose production. Using the genetic standard line *G. hirsutum* cv. TM-1 (Kohel *et al*., 1970) grown under light and temperature-controlled conditions, we sampled daily from 6 to 24 days post anthesis (DPA) to evaluate changes in gene expression during key stages defining the qualities of mature cotton fiber. We characterize gene expression patterns in the context of a developmental time series and use multiple methods to understand the relationships among genes, finding that gene expression is highly coordinated with over half of expressed genes gradually increasing or decreasing in expression throughout the time period studied. We also note a major transcriptomic shift corresponding to the start of the transition phase (Figure 1) and use network analyses to determine putative relationships among key genes. We combine gene expression data with proteomic, glycomic, and phenotypic surveys in the same accession (*G. hirsutum* cv. TM-1) grown under the same conditions and sampled at the same time points to further increase our understanding of the phenotypic consequences of transcriptomic changes. Key candidate genes for control of fiber development are identified and discussed.

## Results

### General description of the data

Gene expression during fiber development was surveyed from the early stages of PCW synthesis through the initiation and maintenance of SCW synthesis (i.e., 6 - 25 DPA; Figure 1). Three replicates were collected for each stage; however, library construction failed for four samples (one each for 20 and 24 DPA and two for 25 DPA). Repeated attempts to generate these replicates were unsuccessful, and thus they were omitted. From the 56 successful samples, we recovered between 1.5 and 264.6 million (M) reads (mean = 41.2 M, median = 36.4 M) per sample. Clean reads were mapped to the 74,776 reference genes, resulting in an average of 55,008 genes expressed at any given time point (Supplementary Figure 1; Supplementary Table 1).

Notably the number of expressed genes (∼74% of transcriptome) is generally stable across replicates and DPA, with the exception of 14 DPA replicate A and the sole 25 DPA replicate (Supplementary Figure 1). Because 14 DPA replicate A had substantially fewer expressed genes than the other replicates, we removed this sample as a potentially early-aborted capsule. We also removed the single 25 DPA sample noted above from subsequent analyses of differential gene expression (DGE) and gene regulatory and/or coexpression reconstruction.

Principal component analysis (PCA) of the expressed genes was used to explore patterns in the data (Figure 2). In general, the first axis (PC1; 56% variance) clustered replicates and sequentially separated DPA along a temporal axis (from left to right). A small gap on the primary axis is observed between 9 and 10 DPA, which reflects the middle of elongation via PCW synthesis. Notably, the largest gap in the primary axis (PC1) is between 16 and 17 DPA, which is at the beginning of the transition stage (∼16-20 DPA; (Tuttle *et al*., 2015)). Interestingly, the initial four timepoints (6-9 DPA) and last six timepoints (19-24 DPA) surveyed exhibited little differentiation along the primary axis, perhaps suggesting relative consistency in expression and/or tighter regulation of expression during the stages of early elongation and early CW thickening, respectively. The seven intervening timepoints (10-16 DPA), which are spread out along the primary axis, are correlated with the majority of elongation before the transition phase begins.

**Figure 2.**
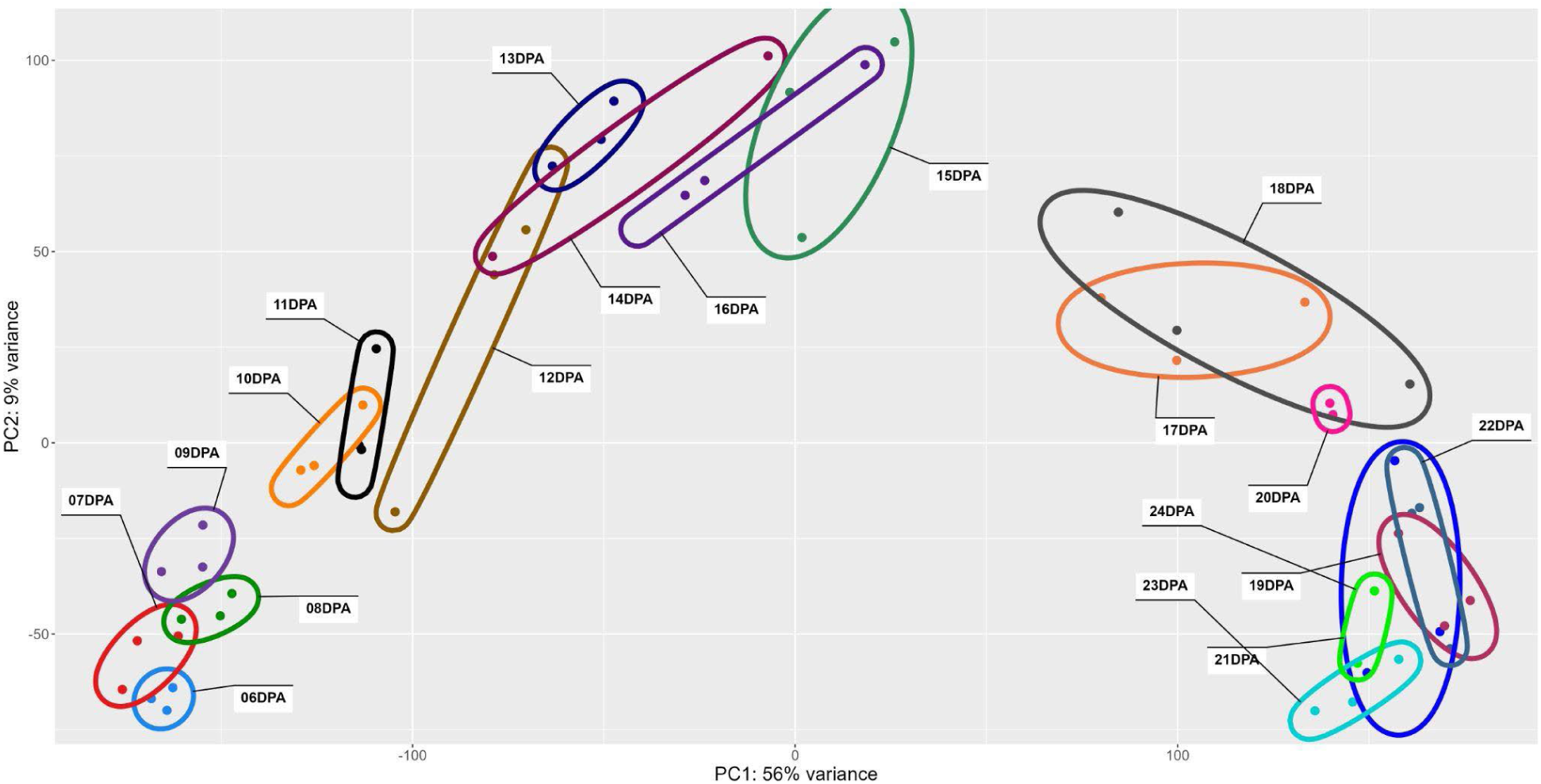
PCA of expression data for cotton fiber sampled daily between 6 and 24 DPA. Each DPA is individually colored and listed, and ellipses encompass replicates for each DPA. First and second axes are displayed, accounting for 54% and 9% of the variance, respectively. PC1 generally separates samples by time, whereas PC2 likely reflects variation between plants or bolls.

### Gene expression trends across fiber development

Differential gene expression was evaluated for all 74,776 reference genes for adjacent stages, as summarized in Figure 2. In general, the number of differentially expressed genes was equivalent between both subgenomes (*i.e.,* AT and DT). Consistent with the aforementioned observation of a distinct difference between 16 and 17 DPA samples (Figure 2), the number of differentially expressed genes (DEG) between those timepoints was more than an order of magnitude greater than most other comparisons (11,417 DEG, versus 16 - 5,562 in other comparison; median = 269 DEG, mean = 1,531 DEG; Supplementary Table 2), suggesting massive changes in gene expression correlated with entering the transition stage. Other, smaller spikes in DEG number were also apparent in the subsequent two comparisons (*i.e.,* 18 versus 17 DPA and 19 versus 18 DPA), as well as between 22 and 23 DPA (Figure 3). Few sharp increases were seen prior to the transition phase, save for small increases in DEG between 7 - 8 DPA and between 12 - 13 DPA. Interestingly, despite the disjunction between 9 and 10 DPA evident in the PCA plot, few genes exhibited significant differences in expression, suggesting that this apparent disjunction between these two DPA is the result of numerous subtle (i.e., not statistically significant) changes in gene expression.

**Figure 3.**
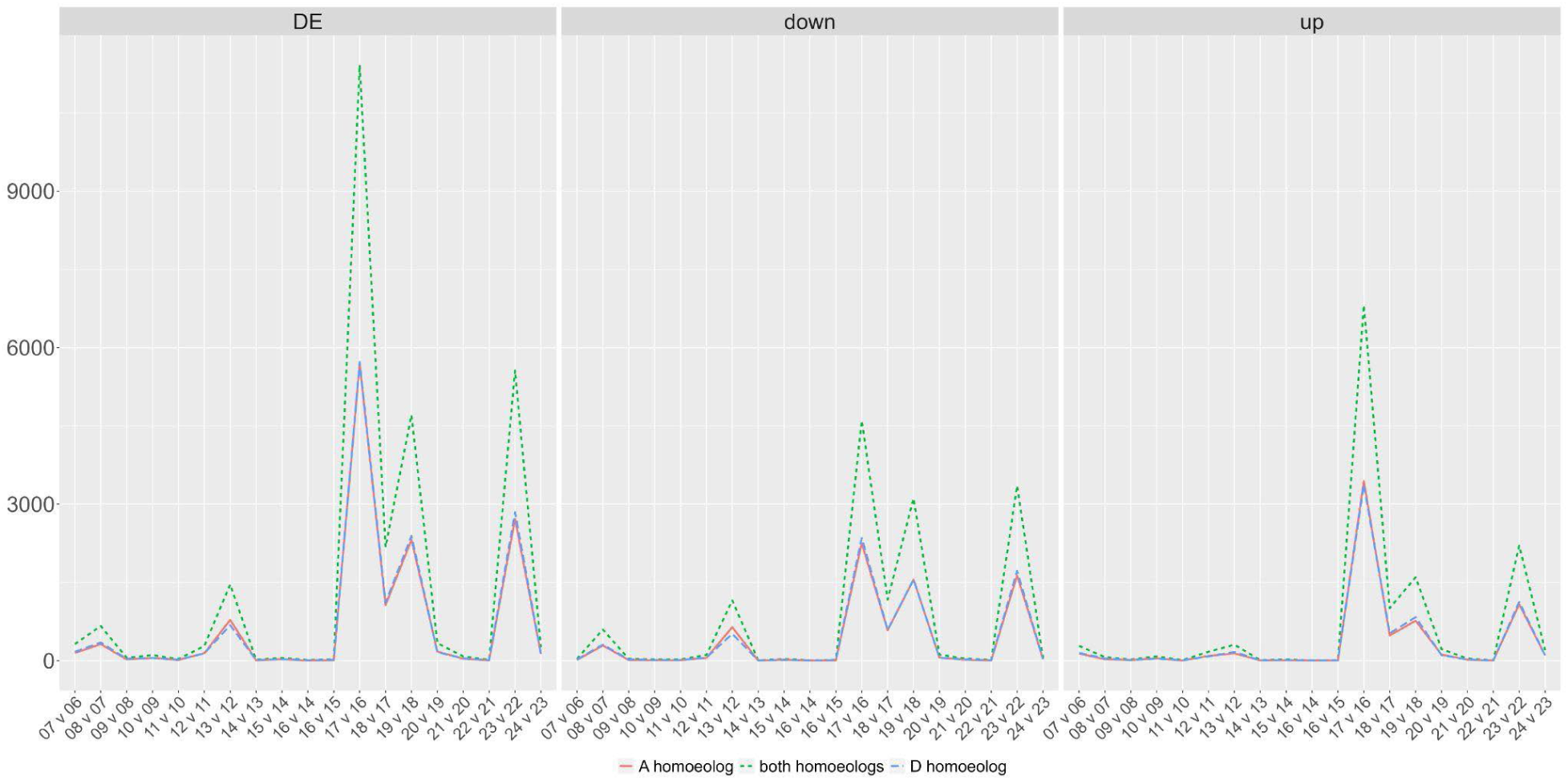
Differential gene expression between adjacent DPA. The number of differentially expressed genes between adjacent DPA comparisons is depicted for the time series. The left panel represents all differentially expressed genes, whereas the middle and right panels are parsed as genes that are up- or down-regulated in the later DPA, respectively. Colors and line types represent either the number of DEG when considering both homoeologs together (green, short dash), the A-homoeolog only (red, solid line), or the D-homoeolog only (blue, long dash).

On average, the number of genes exhibiting down-regulation on adjacent days slightly out-numbered up-regulation (average of 805 versus 726, respectively) across the developmental timeline surveyed here. In nearly two-thirds of the adjacent DPA contrasts (60%; 11 contrasts), the number of down-regulated DEGs at the later days outnumbered the number of up-regulated DEGs; however, the opposite it true when evaluating patterns of differential expression in the context of a timeseries. When fit to a continuous model of gene-wise expression using ImpulseDE2, the number of genes that transition up (Tr-Up; 19,706 genes) or are transiently upregulated (Im-Up; 3,402 genes) during this developmental period (6 to 24 DPA) outnumbers those that transition down (Tr-Down; 14,491genes) or are transiently downregulated (Im-Down; 1,871 genes; Figure 4). The broad classifications of genes in these categories are available in (Supplementary Figure 1). As defined by ImpulseDE2, genes in the transition categories either continuously increase (TrGene-Up) or decrease (TrGene-Down) their expression throughout the sampled time period. The 19,076 genes in the TrGene-Up category encode: glycoside hydrolases with a predicted role in deconstructing CW matrix polymers such as those found in the CFML (Singh *et al*., 2009); transcriptional regulators of SCW synthesis; polysaccharide synthases, including cellulose synthases in all six major classes; accessory protein participants in cellulose synthesis; modulators of the microtubule and actin cytoskeleton; FASCICLIN-like arabinogalactan proteins; hormone response regulators (e.g auxin, brassinosteroid, ethylene, gibberellin, and jasmonic acid); producers and scavengers of reactive oxygen species; and many other proteins that can be logically associated with cotton fiber development (see other text and references in this article). The TrGene-Up category also includes homologs of many other regulatory and structural proteins that have been characterized in cotton or other species (primarily *Arabidopsis*), as well as many uncharacterized proteins.

**Figure 4.**
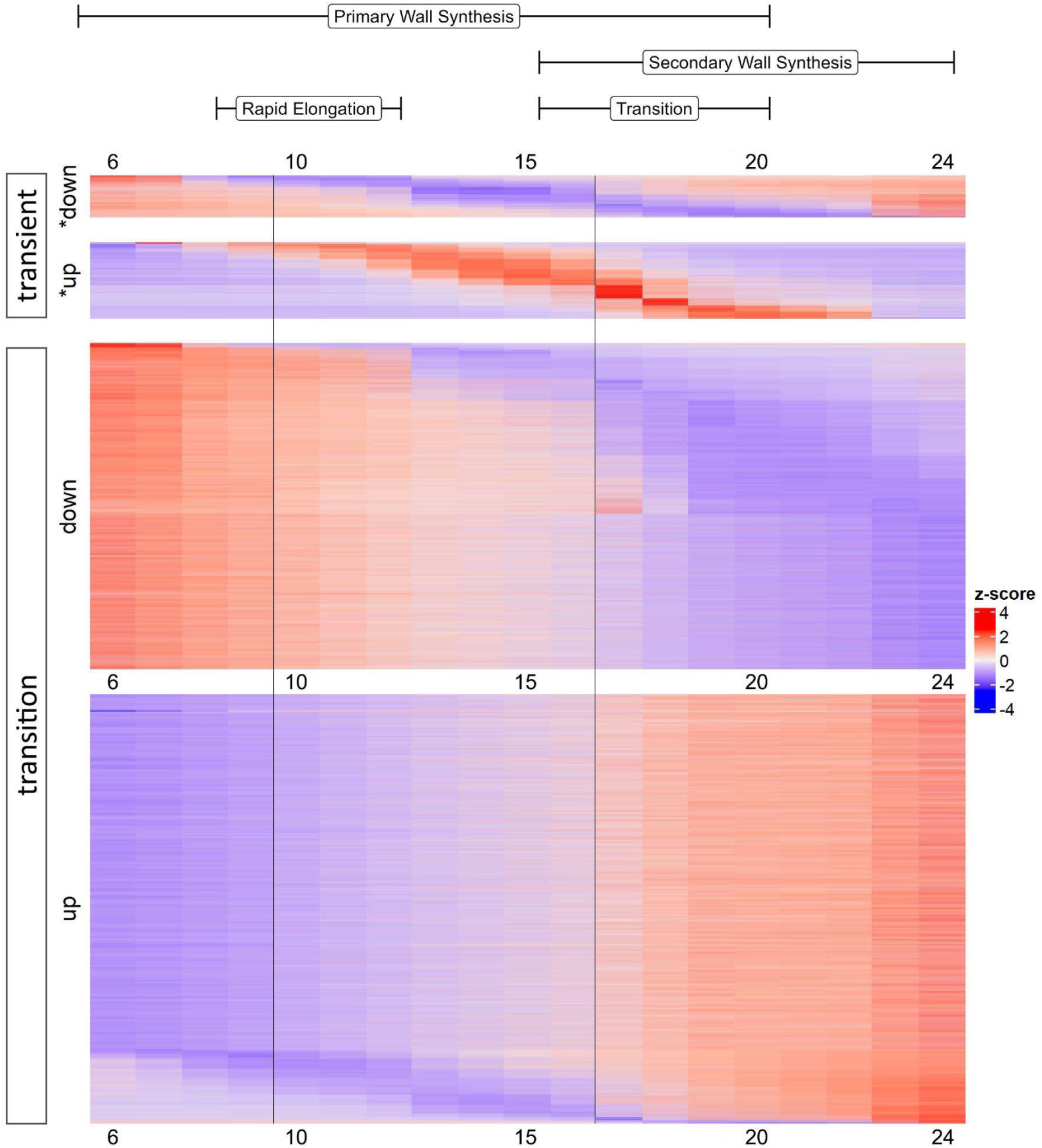
ImpulseDE2 profiles for developing cotton fibers (6 DPA through 24 DPA). Categories include genes whose expression transition up (up, Tr-Up); transition down (down; Tr-Down); impulse up (*up, Im-Up); or impulse down (*down; Im-Down). Colors reflect relative expression levels, where blue indicates lower expression and red indicates higher expression. Bars at the top of the diagram indicate the phase in fiber development covered by those DPA, i.e., PCW synthesis to support rapid elongation, transitional CW remodeling, and SCW synthesis. Vertical black lines indicate the 9-10 and 16-17 DPA gaps from the PCA that also exhibit the most adjacent DPA expression changes.

The transient (or impulse) categories refer to genes whose expression profiles exhibit either increased (Im-Up) or decreased (Im-Down) expression during the middle of the time course and relatively lower or higher expression at the beginning and end, respectively. Interestingly, the beginning of the impulse periods (i.e., where Im-Up and Im-Down genes change expression) coincides with the small disjunction on the PCA between 9 and 10 DPA and the apex of the impulse period coincides with the major shift in gene expression between 16 and 17 DPA. The latter apex is particularly interesting, as it may reflect genes which regulate or participate in the massive changes in gene expression observed at the onset of the transition phase. Gene ontology (GO) analyses of these categories reveal many terms enriched within the Im-Up category for both Molecular Function (MF) and Biological Process (BP), and comparatively fewer terms for the Im-Down category (Supplementary Figures 1 and 2). Among the Im-Up genes (i.e., *up; Figure 4) are genes related to CW extensibility (Jareczek, Grover and Wendel, 2023), which is required for rapid elongation. Interestingly, the proportion of transcription factors in the Im-Up category (10.1%) is significantly lower than found in the Tr-Up category (20.3%, p<0.01 1-sample proportion test) and the proportion of Im-Down transcription factors (17.9%) is significantly greater than that found in the Tr-Down category (10.7%, p<0.01 1-sample proportion test).

Interestingly, the time points sampled captured a small number of genes whose expression increased sharply between 23 and 24 DPA. DEG analysis revealed 201 genes upregulated at 24 DPA relative to 23 DPA (log2 fold change of 0.80 - 33.34), with 82% of the genes having log2 fold change ≥ 2.0. Among these include genes that may be involved in the dominant process of cellulose deposition (see discussion) that begins circa 24 to 25 DPA in cotton fiber, including a GTPase protein (Gorai.011G031400, both ahomoeologs), two NAC transcription factors (Gorai.006G205300.A and Gorai.003G077700.D), and a MYB-like transcription factor (Gorai.001G138800.D).

### Construction of a gene coexpression network

Expression relationships among genes were first analyzed using coexpression network analysis, which places genes into modules based on their correlated expression patterns and summarizes the expression of the genes within each module as the eigengene (i.e., the first principal component of the module).

Approximately 7% (5,237) of the 74,446 genes were removed due to zero variance across the sampled times. The remaining 69,209 genes were placed in 18 modules, referred to as ME0 through ME17 (Figure 5; Supplementary Figure 3), where ME0 (9,748, 14.1%) comprises genes whose expression could not be assigned to a coexpression module (Langfelder and Horvath, 2008). Interestingly, the first two true modules (i.e., ME1 and ME2) each contain over 25% of the genes in the network. ME1 comprises 22,583 genes (32.6%) and exhibits an eigengene profile consistent with increased expression over time (Figure 5; Supplementary Figure 3). Intersection between ME1 and the Tr-Up category of differential expression (above) reveals 16,705 genes from ME1 are also contained within that category (Supplementary Table 3), representing 87.6% of the Tr-Up genes and 74.0% of ME1 genes. Complementing ME1, ME2 (18,919 genes; 27.3% of network) exhibits an eigengene profile consistent with decreasing expression over the time series. Similar to ME1, a majority of Tr-Down genes (12,991 genes, or 89.7%; Supplementary Table 3) are contained within ME2, comprising 68.7% of the total genes in ME2. Notably, the expression profiles of the eigengenes for these first two modules exhibits an axial flip between 16 and 17 DPA (Supplementary Figure 3), reflecting both the disjunction observed in the PCA and the major shift in gene expression exhibited in the time series differential expression analysis.

**Figure 5.**
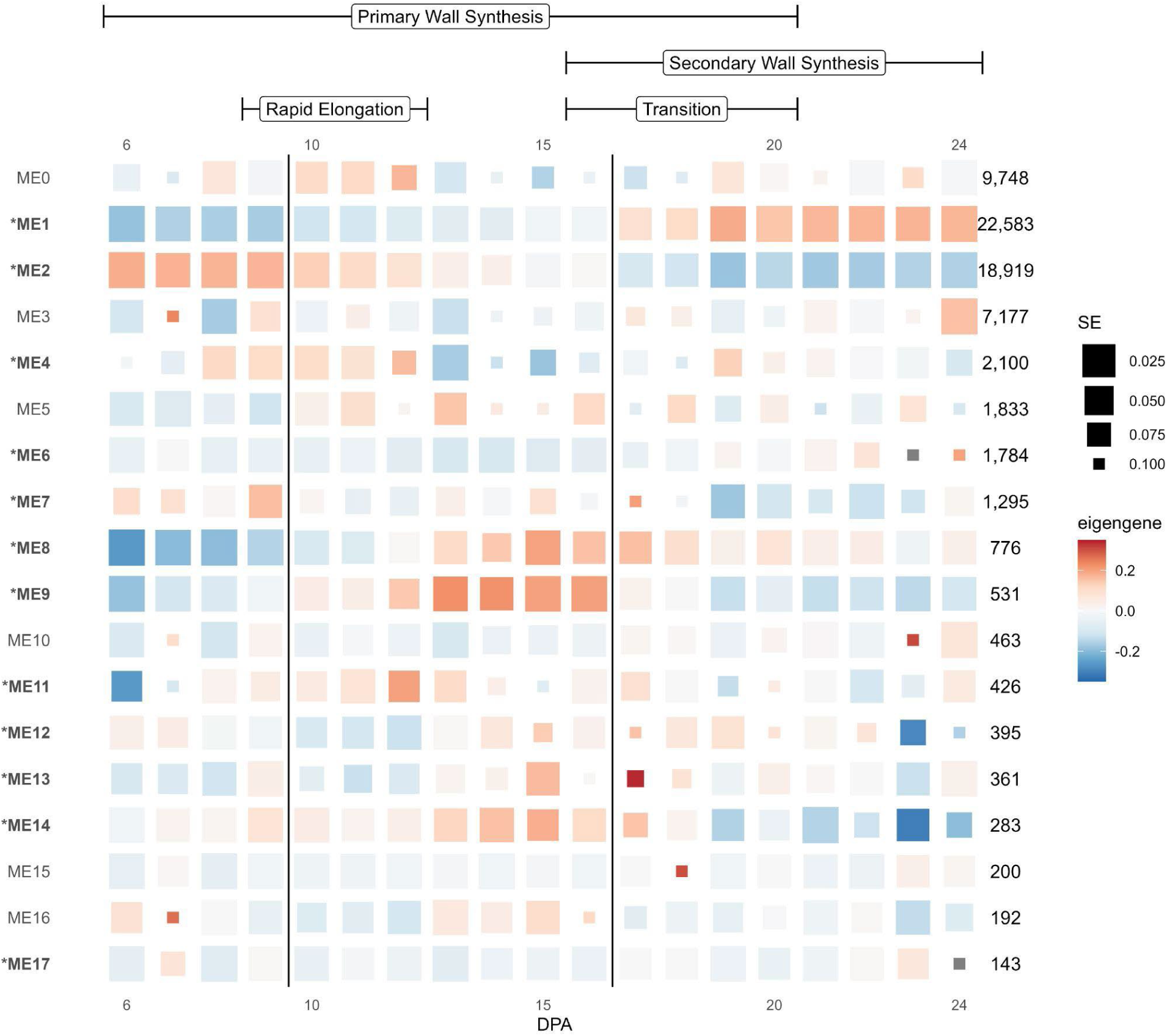
Eigengene expression for coexpression modules derived from cotton fibers developing in stages, as indicated at the top. Modules are listed in numerical order, and modules significantly associated with development are noted with and * and in bold. Colors represent the relative module eigengene expression, and box size represents the standard error (SE), where larger boxes represent eigengene expression values with low SE. Fiber development stages are noted at the top, and the division between 9 versus 10 and 16 versus 17 DPA are noted by vertical lines.

Both ME1 and ME2 also contain relatively high proportions of the Im-Up and Im-Down genes (Supplementary Table 3). ME1 contains 26.4% (899 out of 3,402) of the Im-Up genes and 21.4% of the Im-Down genes (400 out of 1,871), while ME2 contains 10.9% (370 genes) and 42.7% (798 genes), respectively. Although this represents 37.3 - 64.0% of genes contained within each impulse category, these genes represent only about 2 - 4% of the total genes in each module (Supplementary Table 3).

While the expression trajectories of these transiently expressed/suppressed genes may not directly correspond to the module eigengene expression trajectory, their inclusion in these modules may indicate their participation in the general increase or decrease in expression of these modules.

The remaining modules (ME3 - ME17) contain far fewer genes (7,177 to 143, respectively), of which 12 modules are significant with respect to development (p<0.05; anova ME∼sample; Figure 5). Notable among these are ME8 (766 genes) and ME9 (531 genes), both of which contain a relatively high proportion of the Im-Up genes (∼13% each) relative to the remaining modules (except ME1; Supplementary Table 3). In both modules, more than half of the genes are assigned to the Im-Up category (ME8: 450 genes, 58.0% and ME9: 443 genes, 83.4%), which is reflected in their eigengenes, which start with low expression, peak in the middle of the timeseries, and then exhibit declining expression in the later time points; no Im-Down genes are detected in this category. This pattern is particularly apparent in ME9, which exhibits a sharp increase in expression between 9 - 12 DPA and a sharp decline between 16 - 19 DPA, and notably coincides with the fiber developmental periods encompassing rapid elongation and attenuation of elongation, respectively.

Three additional, consecutive modules (i.e., ME12-14) exhibit a high proportion of genes that are considered Im-Up or Im-Down (Supplementary Table 3), which is also somewhat consistent with their eigengene profiles (Figure 5; Supplementary Figure 3). Of those three, ME13 and ME14 have the greatest number of genes in the model that are Im-Up (i.e., 48.2% and 49.8% of module genes, versus 33.9% in ME12), and contain no genes that are considered Im-Down (as was observed for ME8 and ME9).

Conversely, ME12 contains proportionately fewer Im-Up genes along with a small number of Im-Down genes (24; 6.1% of genes in module); however, the eigengene trajectory in ME13 is more similar to ME12 than it is to ME14. That is, both ME12 and ME13 exhibit an increase in eigengene expression starting around 13 DPA that subsequently plummets at ∼23 DPA. ME14 exhibits a dissimilar profile (i.e., increasing steadily from 6 DPA followed by a sharp decline at 19 DPA) to both of these, suggesting that it may reflect a different aspect of fiber development.

With respect to the remainder of the Im-Down category genes, fewer modules (aside from ME1 and ME2) exhibit a relatively high number of these genes relative to the abundance in other modules (Supplementary Table 3). Interestingly, ME0 (i.e., unplaced genes) contains the third greatest number of Im-Down genes after ME1/ME2, perhaps indicating a role for some of these genes that is unclear from the current coexpression analysis. After ME0, ME4 and ME6 contain the most genes from the Im-Down category (ME4=157 and ME6=114), comprising 7.5% and 6.4% of the genes contained within each module, respectively. The eigengene for ME4 (Figure 5, Supplementary Figure 3) exhibits a transient-like pattern of expression, exhibiting a marked reduction between 13 and 18 DPA after which it sharply increases before tapering to 24 DPA. ME6, on the other hand, exhibits low expression until about 22 DPA, where it displays a sharp peak between 22 and 24 DPA, potentially indicating genes important for SCW synthesis, although the standard error for these DPA is high. Nevertheless, 243 genes from ME6 also exhibit significant DE between 22 and 23 DPA, most of which are classified as Tr-Up (226 genes).

GO annotations for these genes are diverse, relating to metabolic processes (e.g., lipid, carbohydrate, and cellular), stimulus/stress response, etc.

### Correlations between coexpression modules and measured phenotypes

We correlated module eigengenes with phenotypic data gathered from the same accession (i.e., *G. hirsutum* cv TM-1) across the same developmental period (Figure 6; Supplementary Table 4; (Swaminathan *et al*., 2024); Howell et al, in prep). As expected from the large number of genes present in the first two modules (22,583 and 18,919 genes, respectively) and the highly canalized nature of fiber development, most traits were significantly correlated (or inversely correlated) with those modules. Those molecules that contribute to CW development (e.g., encode genes involved in pectin, hemicellulose, and cellulose biosynthesis; (Swaminathan *et al*., 2024)) were strongly positively correlated with ME1, which increases in expression over development and strongly negatively correlated with ME2, which decreases over time (Figure 6). Likewise, fiber length (Howell et al, in prep) was strongly positively correlated with ME1 and negatively with ME2; however, these two traits also exhibit relatively strong, significant correlation with ME8 as well. As expected by the enrichment of Im-Up genes in this module, ME8 expression is impulse-like (Supplementary Figure 3), whereby expression starts low, peaks at around 15 DPA, and then decreases again. GO analysis of the 776 genes in this module reveals glycosyl hydrolases, oxidoreductases, and peroxidases (Supplementary Figure 4), which are all important for elongation.

**Figure 6.**
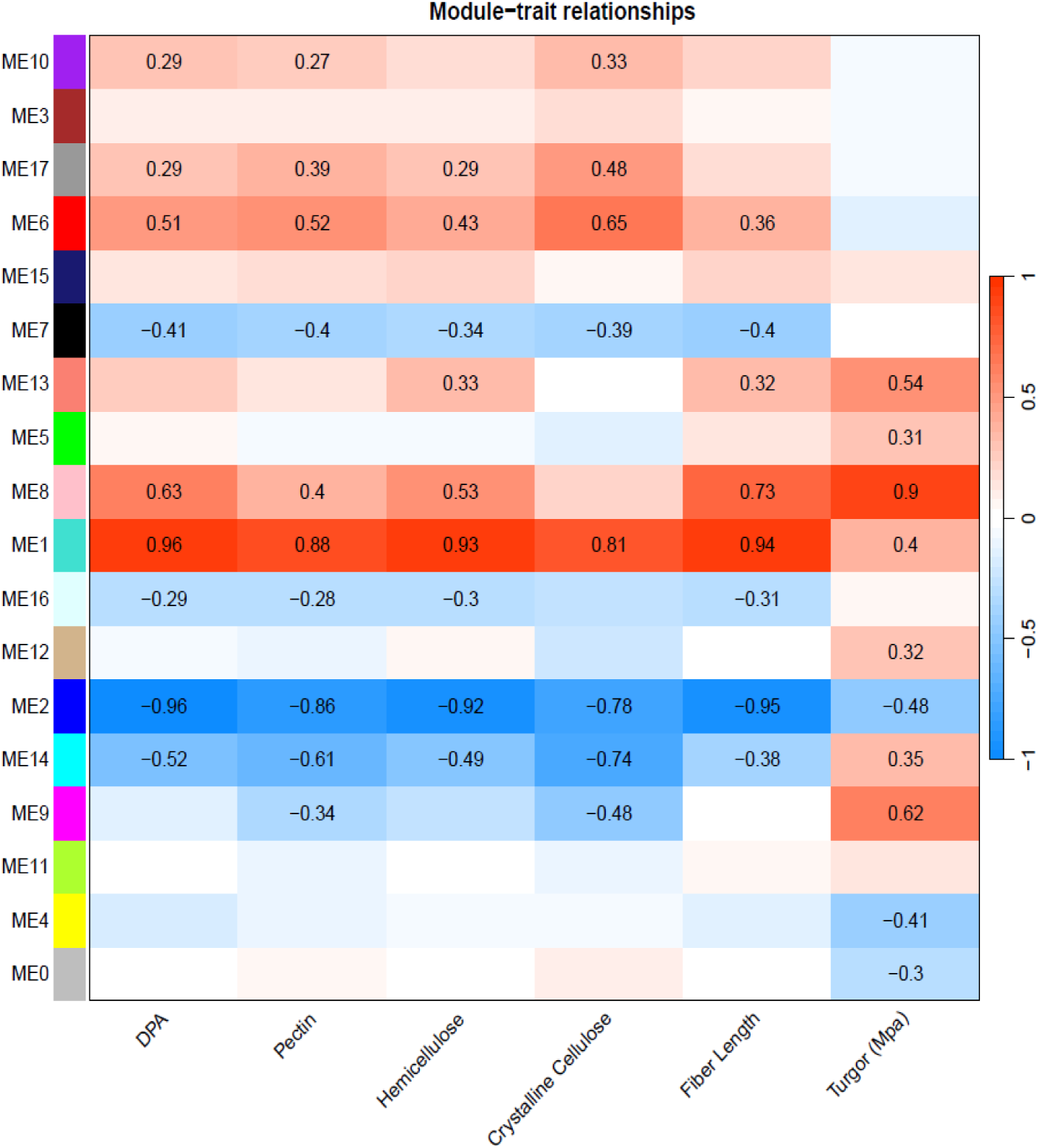
Associations between coexpression modules and phenotypes. Modules are listed on the left, and phenotypes are listed at the bottom. Pectin, hemicellulose, and crystalline cellulose are measured as mg per boll, as per Swaminathan et al (2024), and fiber length is measured as mm, as per Howell et al (in prep). Turgor is interpolated from Ruan et al. (2001), as described in the methods. Positive (red) and negative (blue) correlations are noted, and significant correlations are listed in each box.

Interestingly, turgor pressure exhibited strong correlations with different modules than the rest of the traits. Ruan and coworkers (Ruan et al 2001) used experimental data to estimate turgor values in 5, 10, 16, 20, and 30 DPA, which represented early, mid-, and late-elongation (5 – 16 DPA); transition or early SCW synthesis (20 DPA); and mid-SCW synthesis (30 DPA). For precise correlations with our daily transcriptome data, we interpolated the data to cover 6 – 24 DPA, which showed a gradual increase from 6 – 16 DPA (Supplementary Table 4). Although the interpolated data may be overly smoothed, there was a gradual increase to the peak at 16 DPA (0.67 MPa), followed by a decline through 20 DPA (0.28 mPa) and sustaining of similar values thereafter. Although turgor pressure is somewhat positively correlated with ME1 (r^2^=0.4) and negatively correlated with ME2 (r^2^=-0.48), stronger correlations were seen for ME8 (r^2^=0.9), followed by ME9 (r^2^=0.62) and ME13 (r^2^=0.54). Like ME8, ME9 and (to a lesser degree) ME13 exhibits impulse-like behavior, peaking between 13-16 DPA for ME9 and at 17 DPA for ME13.

### Construction of a crowd network

Because gene network inference algorithms are known to exhibit biases (Marbach *et al*., 2012), we used Seidr (Schiffthaler *et al*., 2023) to generate a crowd network employing 13 algorithms (see methods), including the high performing <u>GE</u>ne <u>N</u>etwork Inference with <u>E</u>nsemble of trees (GENIE3; (Huynh-Thu *et al*., 2010; Greenfield *et al*., 2010)) and Weighted Gene Coexpression Network Analysis (WGCNA). This network was aggregated using the inverse rank product (Zhong *et al*., 2014; Schiffthaler *et al*., 2023), resulting in 2.8 billion (B) edges (30%, or 0.85 B, “directed”edges) between all 74,446 nodes (genes) that exhibit variation among timepoints. Among these, 21,227 undirected and 15,996 directed edges connect nodes representing homoeologs. Since this dense network is composed of both “noisy” edges and those that represent core interactions, we calculated the network backbone to retain only those edges that represent the strongest connections for each node (Coscia and Neffke, 2017; Schiffthaler *et al*., 2023).

Employing a 90% confidence interval reduced the number of edges over 500-fold to 5.1 million (M), which was further reduced to 2.2 M under a 95% confidence interval (see methods). Among these 2.2 M edges, the edge direction (i.e., which member of each pair of adjacent genes operates upstream of the other) is known for 721,101 edges (versus 1.5 M undirected edges). Despite the massively duplicated nature of this polyploid network, <1% of surviving edges (10,761) connect homoeologs; however, just over half of those (5,422) of those are considered directed.

We compared these 2.2M backbone edges to the WGCNA-generated coexpression modules by first clustering the edges of the overall graph using two different algorithms, i.e., Louvain and InfoMap, which produced 188 and 1971 clusters, respectively. By overlapping these clusters with the WGCNA modules, we were able to place genes into 6,519 high confidence groups representing genes which are both placed within the same module and cluster using both algorithms. From these 6,519 groups, slightly less than half (3,094; or 47%) contain at least 1 edge (max: 118,932 edges and 2540 nodes), and possibly represent groups of genes that comprise small subdivisions of the broader gene network (Supplementary Table 5). As expected, the three largest clusters are derived from ME1 (1,500-2,540 genes each out of 22,583 genes total); however, the next largest clusters are not derived from ME1 or ME2 (module membership:∼20k genes each) but are rather formed from genes placed in ME4 (1,438 out of 2,100 genes) and ME5 (1,298 out of 1,833 genes), the latter module which is notably not significant with respect to development. ME4, however, exhibits an eigengene profile consistent with Im-Down between 13 to 18 DPA, a pattern also consistent with the relative abundance of genes exhibiting transient down-regulation expression profiles. While the average and median number of genes per group is relatively low (16 and 3, respectively), 73 groups contain more than 100 genes (average = 424 genes; median = 214) connected by at least 113 edges (average = 11,408; median = 1,620).

Because gene regulatory networks provide insight into the regulatory hierarchies among genes, we isolated those 850 M edges representing the directed gene expression network from the broader crowd network for further analysis. From the top 10% of these edges (i.e., 8.5 M edges), few edges (7,330 or 0.09%) link homoeologs, most of which (5,422 or 74%) are retained in the network backbone described above. Louvain and Infomap clustering of these 8.5 M is similar to the above in that Infomap produces far more clusters (626) than Louvain (5); however, this clustering is notable in the small number of Louvain clusters (5), two of which together contain nearly 92% of genes (Louvain cluster 1 = 35,619 genes, or 48%; Louvain cluster 3 = 32,735 genes, or 44%). When the composition of these clusters is merged with each other and the module designations by WGCNA, it results in 2,206 cluster-groups (Louvain-Infomap-WGCNA), approximately one-third the number of cluster-groups in the backbone that includes both directed and undirected edges. These clusters (Supplementary Table 1; Supplementary Table 5) represent the most confident directed associations among genes in this dataset.

### Phenotypic association between cellulose content and gene regulatory networks

Cellulose deposition in plant cells, including cotton fibers, is a tightly coordinated process driven by cellulose synthase complexes (CSC; (Delmer *et al*., 2024)). Because mature cotton fibers are predominantly composed of cellulose, the orientation of cellulose microfibrils and the amount of cellulose deposited in the SCW are major determinants of key fiber properties (e.g., length and strength). As expected from the integral role of cellulose, crystalline cellulose accumulation (as measured in (Swaminathan *et al*., 2024)) is significantly associated with nearly half (8) of the 17 coexpression modules (Figure 6). Also as expected, cellulose accumulation is most strongly positively correlated with ME1 and most strongly negatively correlated with ME2, the two most gene-rich modules in the coexpression network; however, a strong positive correlation (0.65) was found with the 1,784 genes comprising ME6 and a strong negative correlation (-0.74) was found with the 283 genes comprising ME14. ME6 exhibits generally low expression until around 22 DPA, where it increases rapidly. This module (ME6) notably contains two CesA interacting genes (i.e., a KORRIGAN1-like, KOR1, and a COMPANION OF CELLULOSE SYNTHASE3-like, CC, gene; Gorai.003G089600.A and Gorai.005G256100.D, respectively), which are involved in cellulose synthesis (Pedersen *et al*., 2023).

Phylogenetic analysis of the annotated cotton homoeologs with existing cellulose synthase A (*CesA*) homologs from *Populus trichocarpa* (Kim *et al*., 2019; Paterson *et al*., 2012) and other species revealed 24 *G. hirsutum CesA* genes related to PCW and 12 related to SCW (Supplementary Figure 5; Supplementary Table 6). Due to strong conservation of *CesA* families in vascular plants, expression of *CesA* genes can be broadly partitioned into three major isoform classes each that are expressed during PCW (*CesA1, 3, 6 or 6-like*) or SCW synthesis (*CesA4, 7, 8*), assuming the 10-member *CesA* family of *Arabidopsis* as the canonical reference point (Richmond and Somerville, 2000). Genome duplication in *G. hirsutum* has fostered expansion of the expression set for most of the major *CesA* classes, while also resulting in a non-canonical expression pattern during PCW synthesis for *CesA8-A* homologs. We observed that the three canonical PCW *CesA* classes typically maintain relatively even expression throughout, which may correlate with sampling ending early in SCW synthesis. In contrast, representatives of the three major SCW *CesA* gene classes, which co-function during SCW cellulose synthesis, are all expressed at a low-level beginning at 13 DPA followed by increasing expression during the transition stage and the onset of SCW synthesis. In an exception, there is a *decrease* in expression for both homoeologs of CesA8-A, as noted previously (Tuttle *et al*., 2015; MacMillan *et al*., 2017), which may indicate that only the CesA8-B paralog fulfills the canonical role in SCW synthesis at DPA. In most cases (i.e., CesA7-B, CesA8-A, CesA8-B), the maternal and paternal homoeolog expression profiles were similar within gene; the sole outlier (Figure 7), paralog CesA7-A (Gorai.001G04470), exhibited both comparatively reduced expression in the A homoeolog, as well as a delayed increase in expression (+5 DPA) that peaked at the same time as the rest of the SCW paralogs (∼20 DPA).

**Figure 7.**
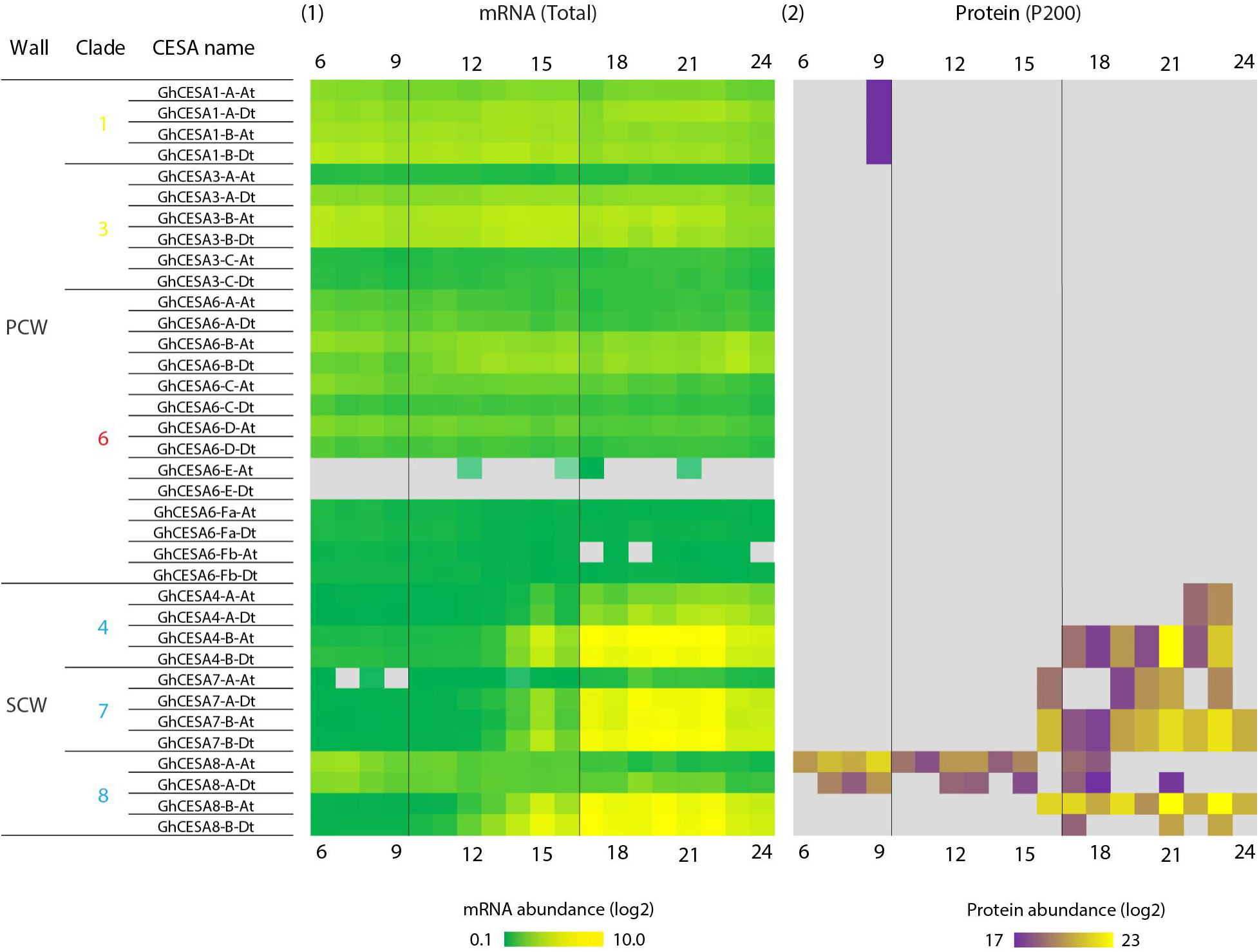
Gene and protein expression for 36 CESA genes across cotton fiber development. CESA homologs that function in primary and secondary wall synthesis (PCW and SCW) are shown. DPA are given across the top and bottom, and key timepoints are noted in vertical black lines. **1.** Gene expression trends for CESA homoeologs for the A and D genomes. **2.** Abundances of CESA proteins isolated from the membrane-associated fraction (P200). Expression for genes and proteins not detected here were rendered with gray background color. The proposed *G. hirsutum* CESA nomenclature and clades are summarized in Supplementary Figure 5 and Supplementary Table 6.

We explored the gene-to-protein expression connection for these genes by comparing the abundance of *CesA* proteins in the membrane-associated (P200) fraction to the transcriptional data for the same DPA profiled here, which detected several secondary wall CESAs from fiber cell extracts (Figure 7). Of the 36 *CesA* homoeologs, all but two (i.e., putative paralogs CesA6-E-At and Dt; see Supplementary Table 1) exhibited measurable *gene* expression (Figure 7) and in all cases both homoeologs were distinguishable in the gene expression data. Due to the challenges of *protein* identification, however, only a subset of those genes were quantifiable via mass spectrometry (16, typically SCW-related; Figure 7; Supplementary Figure 6), most of which were ambiguous with respect to homoeolog of origin (all but CesA-8A and CesA-8B). All of the quantifiable proteins were derived from the membrane-associated (P200) fraction, which is expected due to the multiple transmembrane domains present in CesAs (Li *et al*., 2014).

Notably, the demonstrated presence of CesA8 proteins during PCW synthesis points to the need for future research to understand their specific function at this time (see (Haigler and Roberts, 2019) for a review of less common potential roles for CesA8 orthologs in other species and tissues).

Overall, protein expression profiles for SCW cellulose synthase subunits were generally consistent with their corresponding gene expression profiles, albeit with approximately a 2-3 day difference in expression peaks (Figure 7; Supplementary Figure 6). Abundance profiles for GhCESA4-B, GhCESA7-A/B, and GhCESA8-B proteins were similar to their respective transcripts (Figure 7), being first detected at ∼16 DPA and exhibiting a 2-3 day lag relative to their transcripts. These preliminary results provide a foundation for further exploration of CesA transcript-protein associations during fiber development.

We further compared the expression among CesA isoforms by considering putative regulatory elements involved in CesA gene expression. Using only the directed edges from the Seidr crowd network, we found putative known transcription factors (Jin *et al*., 2014) for 7 genes (10 homoeologs), representing ∼41% of expressed CesA genes (∼30% of CesA homoeologs; Supplementary Table 7). For 6 of the 10 homoeologs, only one transcription factor was directly connected to that gene (3 each for PCW and SCW synthesis); however, for the remaining 4 homoeologs (3 SCW, 1 PCW), between 2-9 putative transcription factors of varying scores and ranks were directly connected to those genes. For the PCW CesA, putative transcription factors were found for the homoeologs GhCESA3-C-At and GhCESA3-C-Dt, although interestingly by transcription factors from different classes (Myb and ARF, respectively; Supplementary Table 7), both of which function in fiber development (Sun *et al*., 2015; X., Zhang *et al*., 2021). The other two PCW genes (GhCESA3-B-Dt and GhCESA6-B-At) are putatively regulated by DOF (DNA-Binding with One Finger) transcription factors, the latter of which has multiple candidate transcription factors from diverse families (Supplementary Table 7). Slightly more putative regulators were found for the SCW genes, likely because the onset of PCW was not sampled here. A single putative TF regulator was associated with GhCESA4-A-At, GhCESA4-A-Dt, and GhCESA4-B-Dt, i.e., a TALE TF (Gorai.003G156000.D; Supplementary Table 7 (Kay *et al*., 2007; Bürglin, 1997)), that rapidly increases in expression beginning around 10 DPA (Tr-Up). Putative regulators for the other subunits were found only for GhCESA7-B-Dt and GhCESA8-B (both homoeologs), each of which had more than one potential TF, sometimes from diverse families. GhCESA7-B-Dt, for example, was associated with 7 possible regulators, including one GATA, two Myb, three NAC, and one TALE TF, with the strongest association (highest ranked edge) connecting GhCESA7-B-Dt to the Myb Gorai.004G138300.D (Supplementary Table 7). Likewise, GhCesA8-B-Dt was associated with 5 possible regulators, including one Dof, two Myb, and two TALE TF, with the strongest association with the TALE Gorai.004G206600.A (Supplementary Table 7). For GhCesA8-B-At, however, there were only two candidate TF, both of which were from the C2H2 family and one of which (Gorai.008G178000.D) exhibited a stronger association.

To understand the position of the SCW cellulose synthase homologs in the context of the broader gene regulatory network (GRN), we explored a subset of the crowd network enriched for the strongest associations between those cellulose synthases and neighboring genes. This strict filtering criteria (see methods) resulted in three subnetworks, a main subnetwork containing representative homologs for each SCW cellulose synthase isoform (i.e., CesA4, CesA7, CesA8; Figure 7, hereafter SCW subnetwork) and two smaller subnetworks that contained only the GhCesA4-A homoeologs or only GhCesA8-B-Dt, both of which were less strongly connected to the larger subnetwork, given our filtering criteria (Figure 8). The large subnetwork contained 3 CESA7s, 2 CESA4s, and 1 CESA8, consistent with the cofunction of the encoded proteins in SCW cellulose synthesis. Both homoeologs of GhCESA4-B are adjacent and linked in the network, occupying a somewhat central location. Notably, some of the putative cellulose synthase transcription factors mentioned above were not present in this subnetwork, likely due to the limited strength of their connections. As expected, several genes that are closely linked to the SCW CesA genes have been previously noted for their importance to fiber development. For example, a FASCICLIN-like arabinogalactan (FLA) precursor is adjacent to GhCESA8-2-At, as is a KOR1-like protein, both of which have been associated with SCW synthesis, but with unproven specific roles so far (Pedersen *et al*., 2023). Another FLA-like protein is proximal to GhCESA7-B-Dt, as are a pectin-lyase and a O-glycosyl hydrolase (GH17) gene, which likely encode enzymes participating in cleavage of CW polymers.

**Figure 8.**
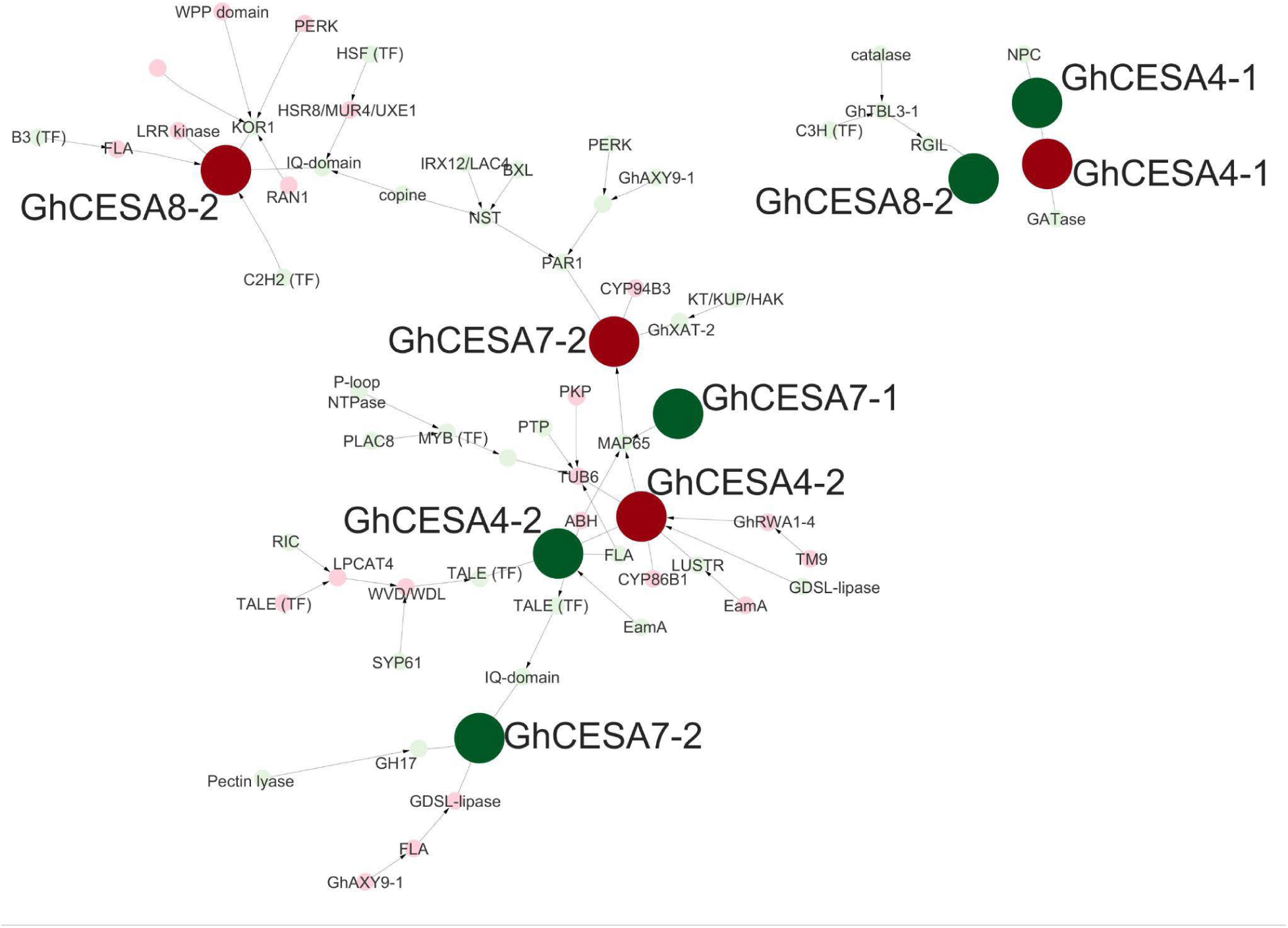
SCW-related CesA subnetwork with neighboring genes. Red circles indicate A homoeologs and green indicate D homoeologs. Further information regarding nodes can be found in Supplementary Table 8 and edge information can be found in Supplementary Table 9. Abbreviations beginning with “Gh” are predicted homologs to the given gene (e.g., “GhCESA4” is homologous to CESA4 from other plants); abbreviations not beginning with “Gh” represent the closest gene annotation, as per (Paterson *et al*., 2012).

Different genes appear adjacent to the A-genome homoeolog for GhCESA7-B, including a gene for xylan side-chain synthesis (GhXAT-2) and a microtubule-associated protein (MAP65-like), the latter of which also appears to be influenced by GhCESA7-A-Dt and both homoeologs of GhCESA4-B. In addition, both GhCESA4-B homoeologs are also linked to previously noted CW genes such as another FLA, a beta-6-tubulin (TUB6), and a reduced wall acetylation gene (GhRWA1-4). Each of these observations has relevance to CW thickening and other transition stage events, as discussed below.

### Phenotypic association between turgor pressure and gene interactions

Although high turgor pressure is implicated in rapid elongation of cotton fibers (Dhindsa *et al*., 1975; Ruan *et al*., 2001; Smart *et al*., 1998), few genes have been identified that may contribute to changes in turgor during fiber development (Sun *et al*., 2019; Ruan *et al*., 2001). Here, we find that turgor is strongly associated with modules that exhibit transient expression patterns, which is perhaps unsurprising given the transient nature of high turgor pressure in driving fiber elongation. Estimated values for turgor pressure were most significantly associated with ME8 (Figure 5) in which gene expression was highest at 15 DPA followed by a gradual decline through 24 DPA. A total of 776 genes are in ME8, including two with functional annotations related to turgor (i.e., a SWEET-like gene (Gorai.003G074400.D) and a PIP-like gene (Gorai.002G198900.D)), both with Im-Up expression patterns similar to the module. There were four genes with functional annotations related to turgor in ME9, which contained 531 genes. ME9 generally contains genes with high expression at 13 - 16 DPA followed by a sharp decline. These ME9 turgor-related genes are: a PIP-like gene (Gorai.006G181300.A), a SWEET-like gene (Gorai.003G074400.A), a SUT/SUC-like gene (Gorai.010G030700.A), and a bHLH transcription factor (Gorai.010G147100.A). As with the two ME8 genes, these genes are considered ImUp, exhibiting increased expression during the intermediate stages and often showing peak expression before 15 DPA when elongation begins to slow down (Figure 9). Notably, the K+ transporter GhKT1 (here, Gorai.012G142000.A and Gorai.012G142000.D) originally noted by Ruan et al (2001) was not found within either of these modules, but rather in ME2 where it exhibits expression that transitions down (considered Tr-Down by ImpulseDE2), congruent with observations in Ruan et al (2001). A different K+ transporter was identified in ME8 (Gorai.009G292800) that was also classified as Tr-Down, and two additional K+ transporters (Gorai.012G082500.A and Gorai.012G082500.D) were identified in ME9, although their expression trend was not described by ImpulseDE2. See discussion for further interpretation of these and other genes relevant to turgor from this module.

**Figure 9.**
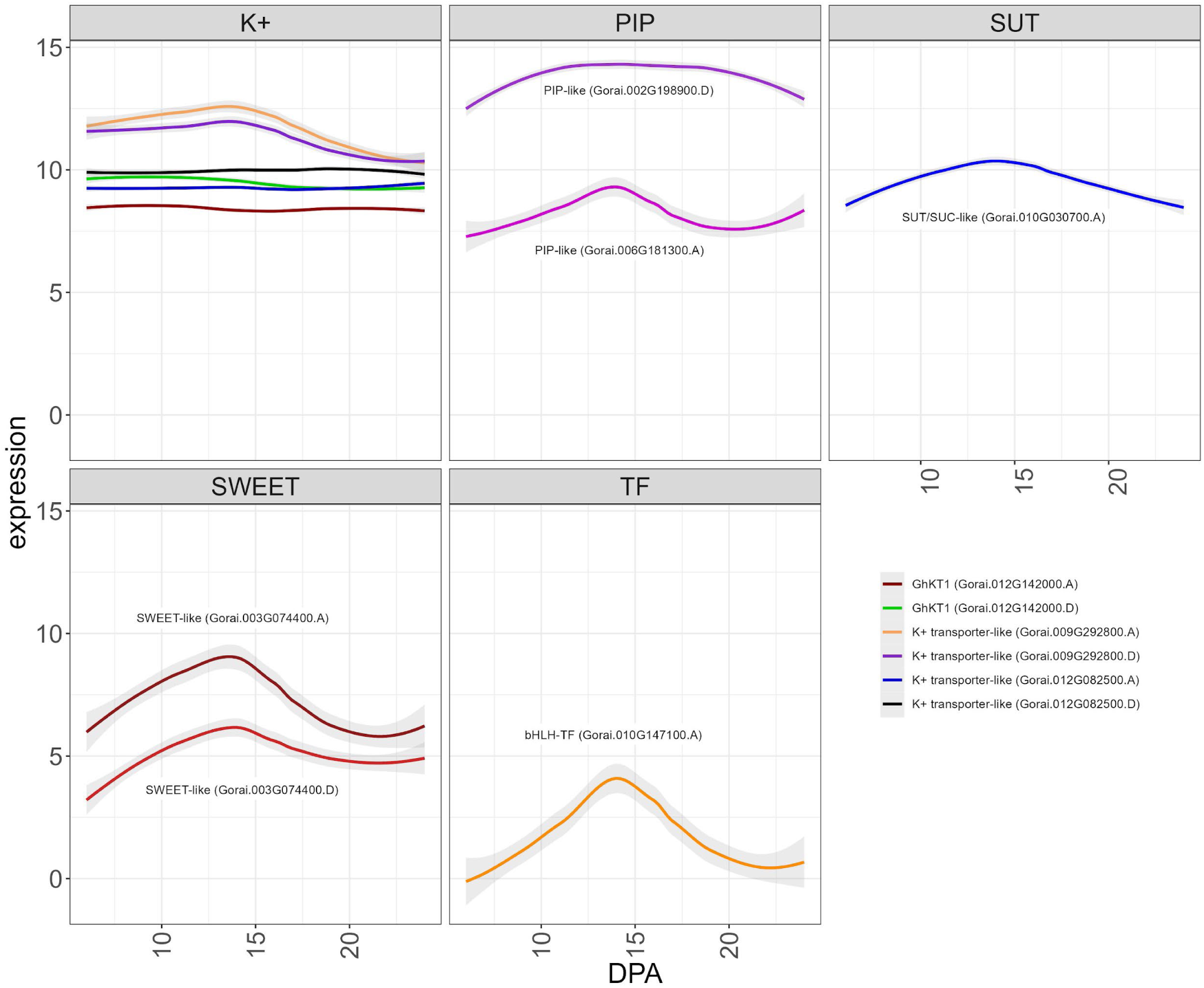
Expression trends for notable genes in ME8 and ME9 with putative relevance to turgor pressure. Genes are partitioned by family, and all lines are labeled except for the potassium transporters (upper left graph), which are distinguished by color. Graphs begin from the initial time point (6 DPA) and continue through the last sampled time point (24 DPA). Intermediate DPA are noted at the bottom of the graphs.

## Discussion

Cotton fiber development entails complex and intricate biological processes encompassing diverse biochemical pathways and transcriptional networks that collectively orchestrate the transformation of newly differentiated fiber initials into mature, elongated fiber cells composed primarily of cellulose. Because of its agronomic importance, understanding the processes that underlie fiber development and how they influence the mature fiber phenotype has been the subject of decades of research. Growth in our understanding of fiber developmental processes has emerged from a great diversity of molecular genetic and genomic studies, ranging from forward genetic analyses of individual genes to large population GWAS studies encompassing multiple accessions. This wealth of prior research has provided a foundation for and motivated the present study, in which carefully controlled conditions were used to constrain experimental and environmental variability. In addition, we used high-dimensionality coexpression and time-series analysis entailing daily sampling of the developing fiber transcriptome to further illuminate the fine-scale molecular basis of fiber development during key stages from early fiber elongation to early CW thickening and associated key fiber modules with important cotton fiber phenotypes.

A striking demonstration of the complexity of cotton fiber development is encapsulated in our observation, initially hinted at over 15 years ago using the less refined technology of the day (Hovav, Udall, *et al*., 2008), that a majority of the ∼70,000 genes (74%) in the cotton genome are expressed in at least one time point in the developing cotton fiber. In general, the transcriptome samples generated here are arrayed along PC1, which divides samples almost linearly according to DPA. Notably, our daily transcriptomic analysis between 6 – 24 DPA diagnosed major, known, aspects of cotton fiber morphogenesis that were hinted at previously but with less temporal resolution. Although prior studies of fiber development in growth chambers or greenhouses have varied in the accession(s) analyzed and the precise growing conditions, there is broad agreement that the transition stage begins at about 14-17 DPA (Avci *et al*., 2013; Tuttle *et al*., 2015; Applequist *et al*., 2001; Chen *et al*., 2012; MacMillan *et al*., 2017). Notably, these same days were among the most dynamic in our analyses (Figures 2 - 4), as indicated by numbers of differentially expressed genes. A genomically global demarcation in gene expression (11,417 DE genes) occurs between 16 – 17 DPA (Figure 2), when the multi-dimensional cellular events characterizing the transition stage are beginning (Haigler *et al*., 2012). Both the number of upregulated and downregulated genes are approximately an order of magnitude greater between 16 and 17 DPA than between any other two adjacent DPA. This impressive and sharp transcriptional demarcation underscores the genome-wide complexity and coregulation of many thousands of genes and their distinctions before and after this transition. Collectively, these data point to this surprisingly brief developmental window as being promising for future insights into the gene regulatory networks and their molecular genetic and chromatin level controls that are key to establishing the SCW synthesis machinery responsible for the development of cotton fiber, and perhaps for its agronomic improvement.

More subtle cellular changes are also revealed by differences in gene expression on adjacent days, noted either by PCA or by adjacent DPA contrasts (Figures 2, 3). A demarcation in gene expression (105 DE genes) occurs between 9 and 10 DPA (see the gap in PC1, Figure 2), when the highest rate of fiber elongation occurs (although the majority of length increase occurs afterwards; (Benedict *et al*., 1999)). At this time, changes also occur in the plasmodesmata that symplastically connect the fiber to the seed.

Specifically, at ∼10 DPA the plasmodesmata become impermeable and structurally begins to switch to a branched form prior to reopening at ∼16 DPA. This change was hypothesized to allow turgor to increase and drive the main phase of fiber elongation (Ruan *et al*., 2001). At the same time, analyzing gene expression changes in the context of a time series revealed expression differences too subtle to be statistically significant in adjacent DPA contrasts. This revealed an interesting difference: although adjacent DPA contrasts (as described above) suggest an overall excess of downregulated genes, the number of genes that increase expression (slowly or rapidly) during the time series is greater than the number of genes that decrease expression. This difference highlights complementary analyses afforded by daily sampling and suggests that expression may increase more slowly, but decline more rapidly, for many of the genes in this key developmental transition. Conceptually, this is consistent with the deposition of nearly pure cellulose into the SCW after the transition stage.

Remarkably, our coexpression analysis partitions nearly 60% of genes into two primary modules reflecting a transcriptionally global synergistic coordination for the singular purpose of fiber CW biosynthesis. These two modules, ME2 and ME1, reflect the major processes of PCW synthesis (to facilitate fiber elongation) and SCW synthesis (to facilitate fiber thickening). Correspondingly, ME2 gene expression generally decreases over time, whereas ME1 gene expression generally increases. ME1 contained the greatest number of genes (22,583) with high expression typically beginning at 17 DPA as CW thickening begins. Conversely, ME2 with the second greatest number of genes (18,919), showed decreasing expression through 17 DPA when elongation was ending. This genome-wide, massive transcriptomic rewiring has few if any precedents in plant biology and begs the question whether other terminally differentiated cell types experience comparable dynamism, or if this property of plant CW development will be discovered to be more common for other cell types.

Although general expression and module association with phenotypes indicates that the fiber transcriptional network is committed to cellulose production during the surveyed timeframe, expression of secondary cell wall CESA genes peaked at around 20 DPA, diminishing shortly thereafter. This result mirrors those from the other cultivated allopolyploid cotton species, *G. barbadense* (Pima cotton), whose developmental timeline is similar albeit with a longer elongation phase (Chen *et al*., 2012; Tuttle *et al*., 2015; Schubert *et al*., 1973). In previous research, gene expression of CW-related genes in *G. barbadense* peaked at 25 DPA (Liu *et al*., 2023), somewhat later than here, although the authors also note that other data demonstrated upregulation of CESA genes at 18 and 28 DPA (Tuttle *et al*., 2015). Given these differences in CESA transcription between species and between studies, it will be of interest to compare the transcriptional program utilized for fiber development in *G. hirsutum* to *G. barbadense* using a similarly controlled and temporally dense sampling of fibers in the latter species as implemented here for the former. This comparison is likely to reveal both commonalities and differences in transcriptional modular deployment, thereby offering possible insight into the important phenotypic traits that distinguish these two important crop species. Likewise, additional sampling is required to further refine the profile of SCW CESA transcription versus translation. At the protein level, SCW CESA subunit production peaks approximately 2 days later, suggesting that post-transcriptional and/or translational control may influence the timing and accumulation of CESA subunits in developing cotton fibers. We note that the longevity of both the mRNA and protein for each SCW CESA isoform was not captured in the present timeline, requiring additional sampling during later timepoints to estimate persistence of each in the cell.

The genes encapsulated by the sharp transcriptional change between the last sampled DPA (i.e., 23 and 24 DPA) also hint at gene expression changes underlying the switch to massive cellulose production. These final sampled DPA correspond to: (a) the highest rate of dry matter accumulation beginning at 24-25 DPA in cotton fiber in this and other studies (Avci *et al*., 2013; Schubert *et al*., 1973); and (b) about 50% (w/w) crystalline cellulose in *G. hirsutum* var TM-1 fiber cell walls by this time, as observed in the current work and previously (Abidi *et al*., 2014). Consistently, spectroscopic analyses show that cotton fiber cellulose begins to exhibit greater self-aggregation around this time (Abidi *et al*., 2014; Lee *et al*., 2015), which is correlated with its progressively increasing proportion in the SCW (Meinert and Delmer, 1977). Genes encoding regulatory proteins that were upregulated in this last surveyed time period were predicted to be positive regulators of mainly cellulose synthesis, which characterizes the final stage of cotton fiber SCW deposition through about 45 DPA.

These last two time points sampled (24-25 DPA) are followed developmentally by streamlined cellulose production in cotton fiber, in which cotton fiber diverges from other plant SCWs to achieve about 95% cellulose content at maturity. This developmental divergence among species is important from fundamental and applied viewpoints; therefore, we highlight genes upregulated at the end of the sampled time series (23 DPA versus 24 DPA) that could logically encode positive regulators of cellulose deposition and be candidates for future research. Two alleles of GhRAC13 (Gorai.011G031400.A and Gorai.010G242900.D; a small, signaling, GTPase protein; see (Didsbury *et al*., 1989) for the meaning of RAC) are upregulated between 23 and 24 DPA, which could result in activation of NADPH oxidase and, consequently, an increasing concentration of H2O2 that stimulates CW thickening (Potikha *et al*., 1999; Delmer *et al*., 1995). NAC transcription factors, all of which contain a conserved N-terminal NAC domain (Aida *et al*., 1997), are also likely to be important. Two NAC alleles (Gorai.006G205300.A and Gorai.003G077700.D) that resemble NST1/SND1 in other species are significantly upregulated at 24 DPA and are able to activate SCW synthesis (Tuttle *et al*., 2015; MacMillan *et al*., 2017). An allele of another high-level SCW transcription factor (Gorai.001G138800.D, resembling MYB83/AT3G08500; see (Ambawat *et al*., 2013) for the meaning of MYB) is also significantly upregulated at this final timepoint. In primary xylem and wood, these transcription factors and others downstream regulate the synthesis of cellulose and other SCW components (Zhong *et al*., 2019; Zhang *et al*., 2018); however, MYB46, an ortholog of the MYB83-type transcription factor upregulated here, can directly bind to the promoters of SCW CESAs and upregulate crystalline cellulose content when over-expressed in *Arabidopsis (Kim et al., 2013)*. We suggest that the upregulation of apparent orthologs of NST1/SND1 and MYB83, along with other direct regulators such as RAC13, may underlie the dominance of cellulose synthesis in cotton fiber after 24 DPA. Notably, putative orthologs of other SCW transcription factors, as inferred from studies of primary and secondary xylem in various species, are expressed in cotton fiber later in developmental time (Tuttle *et al*., 2015; MacMillan *et al*., 2017), which highlights the value of the day-by-day sampling leveraged in this study that captured the first apparent day of transcriptional change to support mainly cellulose synthesis. Further exploration of gene expression changes demarcating these latter DPA, including transcription factors and regulatory genes, and their associations within the GRN underscores the usefulness of this dataset in further exploration of how the synthesis of other typical SCW polymers is downregulated, enhancing our prior insights into how cotton fiber has no or very low lignin (Tuttle *et al*., 2015; MacMillan *et al*., 2017).

### Insights into phenotype via network analysis

Network analysis provides the opportunity to gain insight into the gene relationships that underlie phenotypes. While the spatiotemporal dynamics of several polysaccharides are important for conferring properties relating to fiber quality, we focus here on cellulose accumulation during SCW. The primary GRN that contains representatives of all three main classes of SCW cellulose synthases (CESA4, CESA7, and CESA8; Figure 8) is broadly relevant to events occurring during the transition stage between PCW and SCW synthesis in cotton fiber. Most of the genes in the GRN have increased or sustained expression during the transition stage. Predictions from the function of *Arabidopsis* homologs support the association of known processes with the SCW CESA GRN. Beyond the increased expression of SCW CESAs, an essential gene for cellulose synthesis, KOR1 (Gorai.010G143300.D, AT5G49720.1) is network-adjacent to GhCESA8-2-At. The glucanase-like KOR1 protein interacts with the active cellulose synthase complex (CSC) during cellulose microfibril formation, although its function in vivo is unknown (Delmer *et al*., 2024). Increased cellulose synthesis requires more CSCs to be exported to the plasma membrane, and a phosphoserine protein phosphatases superfamily protein (PAT; Gorai.011G011300.D, AT1G05000.1) can function in this intracellular trafficking (McFarlane *et al*., 2021), as can SYNTAXIN OF PLANTS61 (SYP61; Gorai.009G166000.D, AT1G28490.2) that is able to transport CESAs and KOR1 ((Worden *et al*., 2015) and references therein). Abundant, highly-organized microtubules help to regulate the delivery and function of CSCs in the plasma membrane during SCW formation (Seagull, 1993; Schneider *et al*., 2021). Members of the GRN related to microtubule function include: WAVE-DAMPENED 2-LIKE3 (WDL3 aka WVD/WDL; Gorai.011G171200, At5G61340) (Liu *et al*., 2013), which is involved in the stabilization of cortical microtubules; and MICROTUBULE-ASSOCIATED PROTEIN65-8, which is involved in microtubule bundling during SCW synthesis in tracheary elements (MAP65; Gorai.005G168400.D, AT1G27920.1) (Mao *et al*., 2006). Changes in the microtubule array correlate with an increasingly steep orientation of microtubules and cellulose microfibrils relative to the fiber axis in the distinct ‘winding’ CW layer that is deposited during the transition stage (Meinert and Delmer, 1977; Seagull, 1993). Numerous proteins in the SCW CESA GRN that relate to CW polymer degradation or modification and xylan synthesis are discussed further based on daily characterization of the cotton fiber glycome conducted in parallel to this transcriptomic study (Swaminathan *et al*., 2024). Consistent with the major transcriptional change that occurs at 16 DPA between PCW synthesis (ME2) and SCW synthesis (ME1), the GRN defined by SCW CESAs reflects regulatory processes at several levels including hormones, calcium, management of hydrogen peroxide [a stimulus for the transition to SCW synthesis in cotton fiber (Potikha *et al*., 1999)], protein phosphorylation, transcription factors, sugar and ion transporters, and proposed cell surface glycoprotein sensors (the FLA proteins; see (Pedersen *et al*., 2023)). While it is beyond the scope of this article to discuss all the available functional studies of the genes represented in this GRN, this overview establishes the relevance of the SCW CESA GRN for future research on the control of cotton fiber development and quality.

Turgor pressure, which is regulated through osmotic pressures, is an essential force for plant cell expansion (Zimmermann, 1978; MacRobbie, 2006; Steudle and Zimmermann, 1977). In cotton fibers, high turgor pressure is implicated in rapid elongation (Dhindsa *et al*., 1975; Ruan *et al*., 2001; Smart *et al*., 1998). Turgor pressure is generated by the accumulation of osmotically active solutes like malate (Thaker *et al*., 1999), potassium (Dhindsa *et al*., 1975), and soluble sugars including sucrose (Ruan, 2005) in the central vacuole, followed by the influx of water. During several days within the rapid elongation period, the pressure within the fiber cells increases in association with symplastic isolation as the plasmodesmatal connections to other seed epidermal cells transiently close by the synthesis of callose plugs (Ruan *et al*., 2001). While candidate genes for regulating synthesis and importation of water and solutes have been suggested (Sun *et al*., 2019; Ruan *et al*., 2001), key proteins involved in turgor pressure regulation in cotton fiber remain enigmatic.

In contrast to other fiber phenotypes discussed here that were strongly associated with either ME1 or ME2, the changing turgor pressure estimates derived from prior data (Ruan *et al*., 2001) (Supplementary Table 4) were strongly associated with ME8, which exhibits an impulse-like expression profile for the module eigengene, or with ME9. Both ME8 and ME9 reflect transient gene up-regulation during the latter part of elongation when the plasmodesmata are closed and turgor pressure is increasing. Afterwards, these modules reflect a sharp (ME9) or gradual (ME8) decline in gene expression in the transition stage when fiber elongation is slowing.

Within ME8 or ME9, results implicated genes of four major types as potentially underpinning high turgor in cotton fiber, with cotton and *Arabidopsis* homolog names as follows: a SWEET-like gene (Gorai.003G074400.A and D; At4g10850, AtSWEET7; SUGAR WILL EVENTUALLY BE EXPORTED TRANSPORTER); a SUT/SUC-like gene (Gorai.010G030700.A, At1g09960, SUCROSE TRANSPORTER); two PIP-like genes (Gorai.002G198900.D; Gorai.006G181300.A; At4g35100 PIP2;7; AT4G00430.1, PIP1;4; PLASMA MEMBRANE INTRINSIC PROTEIN); and a bHLH transcription factor (Gorai.010G147100.A; At1g61660, AtBLH112; BASIC-HELIX-LOOP-HELIX). Some members of the sugar transporter families have been characterized in the context of loading photosynthetic sugar into the phloem of *Arabidopsis* leaves, as recently reviewed (Xu and Liesche, 2021). This analogy supports the putative role of the cotton fiber homologs in turgor pressure generation; however, only tentative inferences are appropriate, given evidence that AtSWEET7 functions as a glucose and xylose transporter in engineered yeast (Kuanyshev *et al*., 2021). Characterized sugar transport mechanisms including these protein families often include an apoplastic component (Xu and Liesche, 2021), which would be necessary when cotton fiber plasmodesmata are closed. PIP proteins (like the two detected here), or aquaporins, are well known to transport water across membranes (Jensen *et al*., 2016), and the water will follow an increasing concentration of solutes into the central vacuole to increase turgor pressure. Reduced expression of PIP genes was correlated with shorter mature fibers in transgenic cotton (Li *et al*., 2013) and natural mutants (Naoumkina *et al*., 2015). The AtBLH112 transcription factor acts to increase the synthesis of proline, which is an osmoticum and a free radical scavenger, and to increase the synthesis of enzymes that help to mitigate reactive oxygen stress (Liu *et al*., 2015). Given the role of hydrogen peroxide in triggering the transition stage in cotton fiber (Potikha *et al*., 1999), further research will be needed to determine the role(s) of the cotton homolog of AtBLH112 found in the turgor-associated ME9. In general, the potential role and relevance of these specific genes/proteins to turgor pressure must be functionally tested in cotton itself.

## Conclusions

Here we have characterized the *G. hirsutum* cotton fiber transcriptome with unprecedented daily resolution in plants grown in a growth chamber with uniform light and temperature cycling. The data encompass the 6 – 24 DPA period of fiber development, inclusive of high-rate primary cell elongation, the transition stage to secondary wall synthesis, and thickening of the secondary wall by mainly cellulose deposition. Overall, we report that fiber development involves a dramatically dynamic, genome-wide coordination during which approximately half of the transcriptome increases or decreases expression as development progresses. Our results revealed major gene expression modules associated with known aspects of fiber development, such as the switch from PCW to SCW synthesis. These co-expression modules contain genes, many of which we highlight here, that can be functionally characterized in future research. Sampling at daily intervals also revealed other, more transient gene expression profiles. Some of the transiently expressed genes may prove to be key regulators of important processes, such as turgor pressure, warranting further functional testing. Others may implicate as yet undescribed cellular changes in cotton fiber, stimulating further research. For major discontinuities in gene expression on adjacent days, e.g. 16-17 DPA, even more fine scale temporal sampling will be worthwhile in the future. Applying this approach to other species, e.g. *Gossypium barbadense* with higher fiber quality, or cultivars with different fiber properties, may also be promising directions for studies aimed at understanding evolutionary divergence and crop improvement, respectively. The concurrent proteomic, metabolomic, and phenotypic surveys cited here will provide additional insight into the molecular underpinnings of cotton fiber development and should be generally applicable to the fiber of other modern *G. hirsutum* accessions grown under non-stressful conditions.

## Methods

### Plant growth and sampling

Multiple plants for *Gossypium hirsutum* cultivar TM1 were grown from seed in two gallon pots in growth chambers at Iowa State University (ISU). Growing conditions were standardized on Conviron E15 growth chambers with a relative humidity of 50-70% and a photosynthetic photon flux density (PPFD) of 500 μmol m^−2^ s^−1^. Seeds were sown directly in a soil mixture prepared as 4:2:2:1 soil:perlite:bark:chicken grit. Seeds were germinated and subsequently grown under the same growth chamber conditions, i.e., 16 hour days with 500 umol of light and a temperature of 28°C. A gradual increase in photon intensity was set for the first and last 30 minutes of each day (15 minutes at 166 umol photons + 15 minutes at 336 umol photons). Plants were permitted full dark overnight (8 hours) and growth chambers were cooled to 23°C.

Flowers were hand (self)pollinated using a cotton swab and tagged on the day of anthesis (flowering; 0 DPA). Three samples (replicates) were collected daily during fiber development from 6 DPA (elongation) to 25 DPA (SCW synthesis) for a total of 60 samples (3 replicates x 20 days). Replicates were typically from different plants, aside from two 7 DPA replicates, which were derived from the same plant. Fiber was harvested by extracting whole locules from the bolls prior to flash freezing in liquid nitrogen.

Harvested fiber (in locules) was stored at -80°C until RNA extraction.

### RNA-extraction and RNA-seq

Total RNA was extracted from each sample using a modification of the Sigma Plant Spectrum Total RNA kit (Sigma-Aldrich). First, frozen fibers were ruptured by vortexing locules with ≤106 μm acid-washed glass beads (Sigma-Aldrich) in liquid nitrogen for all DPA, and RNA was extracted using the Spectrum kit including optional washes. The extracted RNA was further purified using phenol-chloroform, as previously described (Hovav, Chaudhary, *et al*., 2008). RNA quality was assessed by the ISU DNA facility using the Agilent 2100 Bioanalyzer, and samples passing quality control (QC) were submitted for RNA-seq at the ISU DNA facility. All three replicates passed QC for each DPA, except for 20 DPA (2 replicates), 24 DPA (2 replicates), and 25 DPA (1 sample only). Although multiple attempts were made to recover additional replicates for these later-stage DPA, these attempts were unsuccessful, due to challenges in extracting RNA from high-cellulose samples. These samples were subsequently omitted, along with a single 14 DPA sample, which exhibited low recovery of gene expression.

Libraries were constructed at the ISU DNA facility using the NEBNext Ultra II RNA Library Prep Kit and sequenced on the Illumina NovaSeq 6000 as paired-end 150-nucleotide reads (PE150). Raw reads were quality and adapter trimmed using trimmomatic version 0.39 (Bolger *et al*., 2014) from Spack (Gamblin *et al*., 2015) as trimmomatic/0.39-da5npsr. Only surviving read-pairs (minimum length of 75nt per read) were retained for expression and network analyses.

### Reference transcriptome generation and mapping

A species-specific, homoeolog-diagnostic reference transcriptome was generated using the *G. raimondii* genome annotation (Paterson *et al*., 2012) in conjunction with species/homoeolog-specific SNP information (Page *et al*., 2013) and a custom script available from https://github.com/Wendellab/TM1fiber. This reference has previously been validated as performing well in the polyploid *G. hirsutum (Hu, Grover, Arick, et al., 2021)* and allowing precise assignment of paired homoeologs. Kallisto v0.46.1 (Bray *et al*., 2016) was used to pseudoalign and quantify transcripts from each sample using ‘kallisto quant’ and processed in parallel using GNU parallel v20220522 (Tange, 2022).

Raw read counts were imported into R/4.2.2 (R Core Team, 2022), and the data were normalized using the variance stabilizing transformation (vst) in DESeq2 v.1.36.0 (Love *et al*., 2014) and the design’∼DPÀ. Principal Component Analysis (PCA) was conducted in DESeq2 using ‘plotPCÀ, and the first two axes were visualized using ggplot2 v3.4.0 (Wickham, 2016). Minimum volume enclosing ellipses were added in ggplot2 using the ggforce v0.4.1 (Pedersen, 2022) Khachiyan-based (Khachiyan, 1996) method ‘+ geom_mark_ellipse()’. Samples irregularly placed on the PCA were noted for follow-up, as they may represent pre-aborted bolls. Of these, only the removed 14 DPA sample exhibiting generally low expression was removed.

RNA-seq quality was also assessed by evaluating generalized expression metrics. Specifically, the number of expressed genes per sample (TPM > 0) was evaluated for consistency among replicates, as were the mean, median, and quantiles (in 10% steps) of these metrics. These metrics were plotted across developmental time using ggplot2, and visual outliers were discarded.

### Differential gene expression

Differential gene expression (DGE) was analyzed in DESeq2 using the design ‘∼DPÀ. Contrasts were conducted between adjacent DPA, and p-values were adjusted (i.e., padj) using the Benjamini-Hochberg correction method (Benjamini and Hochberg, 1995). Differential expression was inferred for any contrast where padj < 0.05. Datatables were generated using tidyverse v1.3.2 (Wickham *et al*., 2019), magrittr v.2.0.3 (Bache and Wickham, n.d.), and data.table v1.14.6 (Dowle *et al*., n.d.). Relevant code is at https://github.com/Wendellab/TM1fiber.

Expression trajectories for genes within the time series were estimated by ImpulseDE2 (Fischer *et al*., 2018) in R/4.2.2. Trajectories were classified by ImpulseDE2 into four categories: consistently increasing (up), consistently decreasing (down), impulse up (up*), and impulse down (down*). For the latter two (impulse) categories, the expression trajectories follow a unimodal pattern where the genes in those categories exhibit transiently high (up*) or low (down*) expression during the time course but return their expression to a level similar to the beginning of the time series.

### Co-expression and GRN analysis

Weighted gene coexpression networks were generated for the 18 remaining timepoints using WGCNA (Langfelder and Horvath, 2008). Raw gene expression values were log-transformed using the ‘rld’ function in WGCNA, and 5327 genes with zero variance were removed, leaving 69,209 genes for coexpression network construction. Soft-thresholding powers were evaluated using the function *pickSoftThreshold* and evaluating powers 1 to 10 and even numbers from 12 to 40, resulting in the selection of power=10. The WGCNA function *blockwiseModules* was used for automatic network construction and module detection using a blocksize that would contain all genes (block=70,000). Module significance relative to the time course was assessed using an ANOVA and p < 0.05. Eigengene values across development were visualized in WGCNA, and modules were functionally assessed using topGO (Alexa and Rahnenfuhrer, 2016). Module-phenotype correlations were computed within WGCNA and visualized using ggplot2. Relevant code is at https://github.com/Wendellab/TM1fiber.

Crowd networks were generated using Seidr v0.14.2 (Schiffthaler *et al*., 2023) and combining networks from 13 algorithms (Supplementary Table 10). All networks were generated within Seidr except WGCNA, which was imported from the above analyses. Networks were combined within Seidr using the inverse rank product (IRP) algorithm (Zhong *et al*., 2014; Schiffthaler *et al*., 2023). This aggregated network was pruned using the backbone function in Seidr, which uses a backboning algorithm (Coscia and Neffke, 2017) to remove edges based on standard deviations from the expected value for that edge. In the present, we used ‘seidr backbone -F 1.64’, which corresponds to retaining edges with p< 0.05. Both the initial aggregate network and the backbone network were clustered using the Louvain (Blondel *et al*., 2008) and InfoMap (Rosvall and Bergstrom, 2008) algorithms from the igraph (v1.4.1) package (Csardi *et al*., 2006). Gene clusters from each algorithm were intersected between themselves and the WGCNA-generated modules to form cluster-groups that are composed of those genes that belong to the same module, Louvain cluster, and InfoMap cluster.

Gene regulatory networks were generated by restricting the output from Seidr to only “directed” edges. Again this was done for the aggregate network and the backbone network, albeit with a more relaxed backbone threshold (‘seidr backbone -F 1.64’, or p < 0.05) to recover more edges from the naturally less dense directed network. These networks were Louvain and InfoMap clustered (as above) and intersected with WGCNA modules to generate directed cluster-groups.

### Transcription factor analysis

Transcription factors for the *G. raimondii* genome were downloaded from the PlantTFDB v 5.0 (Tian *et al*., 2020; Jin *et al*., 2017). Both transcription factor (TF) gene ID and family were retained. Expression profiles for transcription factors were extracted from the broader DESeq2 and ImpulseDE2 analyses (above). TF presence in modules and cluster-groups was derived from the above analyses and recovered using tidyverse v1.3.2 (Wickham *et al*., 2019) in R. With respect to the gene network analyses, two types of networks were considered: (1) TFe, or transcription factor extended, which retained edges when at least one of the two nodes was a transcription factor, and (2) TFr, or transcription factor restricted, which only retained edges when both nodes were transcription factors.

### Protein sequence alignments and phylogenetic analysis

Cellulose synthase (CESA) protein sequences from *Populus trichocarpa* (Kim *et al*., 2019) and several landmark species (Lee and Szymanski, 2021) were downloaded from Phytozome V13 (Goodstein *et al*., 2012). A multiple sequence alignment was generated using Clustal Omega at EMBL-EBI (Madeira *et al*., 2022) with the number of combined iterations set to 5 and setting the distance matrix as output. This distance matrix was used for the correlation analysis between protein sequences and transcript abundances (see below). The phylogenetic tree was built from the alignment generated by Clustal Omega on EMBL-EBI (https://www.ebi.ac.uk/Tools/msa/clustalo/).

### CesA network filtering

To evaluate the local neighborhood of the cellulose synthase (CesA) genes involved in SCW synthesis, we targeted genes that belong to the largest WGCNA coexpression module (ME1), Louvain cluster #6, InfoMap cluster #22 (henceforth 1-6-22), which contained 9 of the 12 SCW CesA genes. Using the top 10% of edges in the crowd network (54,705 nodes and 222,490 edges), we extracted only directed edges that included one of the 947 genes (nodes) from the SCW cluster as either a source or target node, resulting in a network composed of 1279 nodes and 1448 edges. We further restricted our edges to those included in the top 10% of edges for this SCW cluster, resulting in 225 nodes and 145 edges. We imported those edges into Cytoscape v3.10.1 (Shannon *et al*., 2003), where we filtered nodes to retain only those with at least one outgoing edge and all CesA genes. We further reduced the network view to include only the nearest neighbors to the CesA genes by iteratively using “Select > First Neighbors of Selected Nodes” five times.

### Isolation of microsome (P200) fraction

The microsome (P200) fraction was obtained from intact cotton fiber tissue from 6 to 24 DPA (McBride *et al*., 2017). Briefly, apoplastic proteins and extracellular vesicles were removed from the intact ovules (∼200 mg) in one locule by dipping each ovule into 5 mL of microsome isolation buffer (MIB) [50 mM Hepes/KOH (pH 7.5), 250 mM sorbitol, 50 mM KOAc, 2 mM Mg(OAc)2, 1 mM EDTA, 1 mM EGTA, 1 mM dithiothreitol (DTT), 2 mM PMSF and 1% (v/v) protein inhibitor cocktail (160 mg/mL benzamidine-HCl, 100 mg/mL leupeptin, 12 mg/mL phenanthroline, 0.1 mg/mL aprotinin, and 0.1 mg/mL pepstatin A)] with 10 minutes incubation under gentle shaking. The ovules were recovered from the MIB buffer and fiber tissues were isolated from seeds as described previously (Lee and Szymanski, 2021). The fiber tissues were homogenized under cold MIB using a Polytron homogenizer (Brinkmann Instruments) and filtered through 4 layers of cheesecloth pre-soaked in cold MIB. Debris in the filtered homogenate was pelleted at 1,000 x g for 10 min using an Allegra X-30R centrifuge (Beckman Coulter Life Sciences). Microsomes were enriched at 200 k x g for 20 minutes at 4°C using a Beckman Optima Ultracentrifuge with TLA110 rotor (Beckman Coulter Life Sciences) and washed twice with MIB. The final pellet was mixed with 200 μL of 8 Urea and incubated for 1 hour at room temperature to denature proteins from membranes. Undissolved debris was removed by centrifugation at 12,000g for 15 minutes using an Allegra X-30R centrifuge. Three biological replicates were prepared.

### Protein mass spectrometry analysis

LC-MS/MS run and peptide identification/quantification were performed as described previously (Lee and Szymanski, 2021; McBride *et al*., 2017)). Briefly, 50 μg of proteins in the P200 fractions were digested using trypsin and digested peptides were subsequently purified using C18 Micro Spin Columns (74-4601, Harvard Apparatus). For each sample, 1 μg was analyzed by reverse-phase LC-ESI-MS/MS using a Dionex UltiMate 3000 RSLCnano System coupled with the Orbitrap Fusion Lumos Tribrid Mass Spectrometer (Thermo Fisher Scientific Inc.). The Andromeda search engine on MaxQuant (version 1.6.14.0) was used for relative protein abundance quantification and protein identification (Cox et al., 2014; Tyanova, Temu, & Cox, 2016). The search parameters were as follows: (1) the match between runs function was set with a maximum matching time window of 0.7 min as default; (2) only proteins identified by a single unique peptide were selected; (3) the same reference generated for RNAseq was used; (4) label-free quantification was selected; and (5) all other parameters were set as default.

### Cell wall and polysaccharide extraction

Alcohol-insoluble CW and subsequently the pectin, hemicellulose and cellulose polysaccharides were extracted from cotton fiber in triplicate using a modification of previous methods (Avci *et al*., 2013) using the same time points sampled above (i.e., 6 to 24 DPA), as per Swaminathan et al (Swaminathan *et al*., 2024). Each cotton boll (stored at -80°C) was thawed until 28°C, at which point fibers were removed using a scalpel and forceps and subsequently placed in a tube on ice. Harvested fibers were ground thoroughly in liquid nitrogen, and the CW was extracted by using a series of organic solvents (Avci *et al*., 2013). From the CW, non-cellulosic polysaccharides, such as pectin and hemicellulose, were extracted, as previously described (Zabotina *et al*., 2012), using 50 mM CDTA:50 mM ammonium oxalate (1:1) buffer followed by 4M KOH, respectively. The final cellulose pellet (containing a mixture of both amorphous and crystalline celluloses) that remained after the 50 mM CDTA:50 mM ammonium oxalate buffer and the 4M KOH extractions was dried, weighed, and analyzed.

### Turgor gene identification

Turgor pressures over the developmental timeline were estimated by inferring intermediate values based on existing measured values (Ruan *et al*., 2001). These data were originally measured by first determining osmolalities (Ruan *et al*., 1995) and converting to MPa using 2.48 MPa per Osm kg^-1^, and then estimating turgor from the difference in osmotic and water potential. Measured values (Ruan *et al*., 2001) include 0.075 MPa (5 DPA), 0.11 MPa (10 DPA), 0.68 MPa (16 DPA), 0.28 MPa (20 DPA), and 0.25 MPa (30 DPA). These points were used to generate a first order b-spline of 100 datapoints in the 5 to 30 DPA interval. The values at 6 to 24 DPA were used as estimates for turgor pressure variability over the time interval of this study.

Osmolytes involved in increasing turgor were identified from the literature (Kopka *et al*., 1997; Dong *et al*., 2018; Ruan *et al*., 2001; Rhodes and Samaras, 2020). *Arabidopsis thaliana* genes involved in producing or transporting these osmolytes were identified in TAIR (Berardini *et al*., 2015; Cheng *et al*., 2017). Putative cotton homologs were identified using the orthologous groups available on Phytozome v12.1 (Goodstein *et al*., 2012) and were assumed to have similar involvement as in *A. thaliana*.

Candidates from this list of turgor-involved genes that were also present in the turgor-associated modules (ME8 and ME9) were identified. Expression trajectories for those 6 genes were extracted from the log-transformed, normalized dataset used in WGCNA, and then smoothed and plotted in ggplot2. T

## Data and code availability

RNAseq reads are available from the Short Read Archive (SRA) under PRJNA1099209. Code used to analyze the data is available at https://github.com/Wendellab/TM1fiber. The mass spectrometry proteomics data have been deposited to the ProteomeXchange Consortium via the PRIDE (Perez-Riverol *et al*., 2022) partner repository with the dataset identifier PXD051704.

## Authors Contributions

JFW, OZ, DBS, and CEG conceptualized the project. PY engaged in data curation. CEG and YL were involved in formal analysis. Funding was acquired by JFW, OZ, DBS, JX, and DGP. SS, AGL, JJJ, YL, XX, MAA, CEG, AHH, PY, and CHH conducted the investigation. JFW, DBS, ELM, CEG, JX, and OZ administered the project. Resources were provided by JJJ, AGL, ERM, MAA, and DGP. JFW, OZ, DBS, CEG, GH, and DGP engaged in supervision of the project. Visualization was conducted by MAA, CEG, YL, and HR. The original draft was written by CEG and CHH, and all authors were involved in manuscript review.

## Acknowledgements

The authors thank Weixuan Ning and Ehsan Kayal for their helpful discussion. The authors also thank the ResearchIT unit at Iowa State University for computational support. We thank the USDA-ARS (58-6066-0-066, Genomics of Malvaceae) for their financial support.

---

**Supplementary Figure 1.** Molecular function GO enrichment word maps for each category from ImpulseDE2: (A) impulse up, 3402 genes; (B) impulse down, 1871 genes; (C) transition up,19706 genes; and (D) transition down, 14491 genes.

**Supplementary Figure 2.** Biological process word maps for GO enrichment for each category from ImpulseDE2: (A) impulse up, 3402 genes; (B) impulse down, 1871 genes; (C) transition up,19706 genes; and (D) transition down, 14491 genes.

**Supplementary Figure 3.** Relative expression of module eigengenes over developmental time. Each module is listed by number and color, as output by WGCNA. The number of genes in each module is listed, and the significance of the module to the developmental timeline (as determined by ANOVA) is listed.

**Supplementary Figure 4.** Molecular function GO enrichment word map for ME8, 776 genes.

**Supplementary Figure 5.** Phylogenetic analysis of CESA orthologs. CESA protein sequences from *Populus trichocarpa* (Kim et al, 2019) and landmark species (Lee and Szymanski, 2021) were downloaded from Phytozome V13 (Goodstein et al, 2012) for the analysis. Phylogenetic analysis was performed by Clustal Omega (https://www.ebi.ac.uk/Tools/msa/clustalo/).

**Supplementary Figure 6.** Profiles of mRNA and protein abundances of selected CESAs that belong to informative groups at protein level. AtDt suffixes reflect ambiguity with respect to homoeolog identification and Dt indicates homoeolog-specific peptides were identified. PCC: <u>P</u>earson <u>C</u>orrelation <u>C</u>oefficient.

## References

1. Abidi, N., Cabrales, L. and Haigler, C.H. (2014) Changes in the cell wall and cellulose content of developing cotton fibers investigated by FTIR spectroscopy. Carbohydr. Polym., 100, 9–16. Available at: 10.1016/j.carbpol.2013.01.074.

2. Ahmed, M., Shahid, A.A., Din, S.U., et al. (2018) An overview of genetic and hormonal control of cotton fiber development. Pak. J. Bot., 50, 433–443.

3. Aida, M., Ishida, T., Fukaki, H., Fujisawa, H. and Tasaka, M. (1997) Genes involved in organ separation in Arabidopsis: an analysis of the cup-shaped cotyledon mutant. Plant Cell, 9, 841–857. Available at: 10.1105/tpc.9.6.841.

4. Alexa, A. and Rahnenfuhrer, J. (2016) *topGO: Enrichment Analysis for Gene Ontology*,.

5. Ambawat, S., Sharma, P., Yadav, N.R. and Yadav, R.C. (2013) MYB transcription factor genes as regulators for plant responses: an overview. Physiol. Mol. Biol. Plants, 19, 307–321. Available at: 10.1007/s12298-013-0179-1.

6. Ando, A., Kirkbride, R.C., Jones, D.C., Grimwood, J. and Chen, Z.J. (2021) LCM and RNA-seq analyses revealed roles of cell cycle and translational regulation and homoeolog expression bias in cotton fiber cell initiation. BMC Genomics, 22, 309. Available at: 10.1186/s12864-021-07579-1.

7. Applequist, W.L., Cronn, R. and Wendel, J.F. (2001) Comparative development of fiber in wild and cultivated cotton. Evol. Dev., 3, 3–17. Available at: 10.1046/j.1525-142x.2001.00079.x.

8. Avci, U., Pattathil, S., Singh, B., Brown, V.L., Hahn, M.G. and Haigler, C.H. (2013) Cotton fiber cell walls of Gossypium hirsutum and Gossypium barbadense have differences related to loosely-bound xyloglucan. PLoS One, 8, e56315. Available at: 10.1371/journal.pone.0056315.

9. Bache and Wickham magrittr: a forward-pipe operator for R. R package version.

10. Benedict, C.R., Kohel, R.J. and Lewis, H.L. (1999) Cotton Fiber Quality. In W. C. Smith, ed. Cotton: Origin, History, Technology, and Production. New York, NY: John Wiley & Sons, pp. 269–288.

11. Benjamini, Y. and Hochberg, Y. (1995) Controlling the False Discovery Rate: A Practical and Powerful Approach to Multiple Testing. J. R. Stat. Soc. Series B Stat. Methodol., 57, 289–300. Available at: http://www.jstor.org/stable/2346101.

12. Berardini, T.Z., Reiser, L., Li, D., Mezheritsky, Y., Muller, R., Strait, E. and Huala, E. (2015) The Arabidopsis information resource: Making and mining the “gold standard” annotated reference plant genome. Genesis, 53, 474–485. Available at: 10.1002/dvg.22877.

13. Blondel, V.D., Guillaume, J.-L., Lambiotte, R. and Lefebvre, E. (2008) Fast unfolding of communities in large networks. J. Stat. Mech., 2008, P10008. Available at: https://iopscience.iop.org/article/10.1088/1742-5468/2008/10/P10008/meta [Accessed June 21, 2023].

14. Bolger, A.M., Lohse, M. and Usadel, B. (2014) Trimmomatic: a flexible trimmer for Illumina sequence data. Bioinformatics, 30, 2114–2120. Available at: 10.1093/bioinformatics/btu170.

15. Bray, N.L., Pimentel, H., Melsted, P. and Pachter, L. (2016) Erratum: Near-optimal probabilistic RNA-seq quantification. Nat. Biotechnol., 34, 888. Available at: 10.1038/nbt0816-888d.

16. Buchala, A.J. (1999) Noncellulosic carbohydrates in cotton fibers. Cotton fibers-developmental biology, quality improvement, and textile processing. *New York*: *Haworth Press Inc*, 113–136.

17. Bürglin, T.R. (1997) Analysis of TALE superclass homeobox genes (MEIS, PBC, KNOX, Iroquois, TGIF) reveals a novel domain conserved between plants and animals. Nucleic Acids Res., 25, 4173–4180. Available at: 10.1093/nar/25.21.4173.

18. Butterworth, K.M., Adams, D.C., Horner, H.T. and Wendel, J.F. (2009) Initiation and Early Development of Fiber in Wild and Cultivated Cotton. International Journal of Plant Sciences, 170, 561–574. Available at: 10.1086/597817.

19. Cheng, C.-Y., Krishnakumar, V., Chan, A.P., Thibaud-Nissen, F., Schobel, S. and Town, C.D. (2017) Araport11: a complete reannotation of the Arabidopsis thaliana reference genome. Plant J., 89, 789–804. Available at: 10.1111/tpj.13415.

20. Chen, X., Guo, W., Liu, B., Zhang, Y., Song, X., Cheng, Y., Zhang, L. and Zhang, T. (2012) Molecular mechanisms of fiber differential development between G. barbadense and G. hirsutum revealed by genetical genomics. PLoS One, 7, e30056. Available at: 10.1371/journal.pone.0030056.

21. Constable, G., Llewellyn, D., Walford, S.A. and Clement, J.D. (2015) Cotton Breeding for Fiber Quality Improvement. In V. M. V. Cruz and D. A. Dierig, eds. Industrial Crops: Breeding for BioEnergy and Bioproducts. New York, NY: Springer New York, pp. 191–232. Available at: https://doi.org/10.1007/978-1-4939-1447-0_10.

21. Coscia, M. and Neffke, F.M.H. (2017) Network Backboning with Noisy Data. In 2017 IEEE 33rd International Conference on Data Engineering (ICDE). pp. 425–436. Available at: 10.1109/ICDE.2017.100.

22. Csardi, G., Nepusz, T. and Others (2006) The igraph software package for complex network research. InterJournal, complex systems, 1695, 1–9.

23. Delmer, D., Dixon, R.A., Keegstra, K. and Mohnen, D. (2024) The plant cell wall—dynamic, strong, and adaptable—is a natural shapeshifter. Plant Cell, koad325. Available at: https://academic.oup.com/plcell/advance-article/doi/10.1093/plcell/koad325/7596221 [Accessed February 9, 2024].

24. Delmer, D.P., Pear, J.R., Andrawis, A. and Stalker, D.M. (1995) Genes encoding small GTP-binding proteins analogous to mammalian rac are preferentially expressed in developing cotton fibers. Mol. Gen. Genet., 248, 43–51. Available at: 10.1007/BF02456612.

25. Dhindsa, R.S., Beasley, C.A. and Ting, I.P. (1975) Osmoregulation in Cotton Fiber: Accumulation of Potassium and Malate during Growth. Plant Physiol., 56, 394–398. Available at: 10.1104/pp.56.3.394.

26. Didsbury, J., Weber, R.F., Bokoch, G.M., Evans, T. and Snyderman, R. (1989) rac, a novel ras-related family of proteins that are botulinum toxin substrates. J. Biol. Chem., 264, 16378–16382. Available at: https://www.ncbi.nlm.nih.gov/pubmed/2674130.

27. Dong, H., Bai, L., Zhang, Y., et al. (2018) Modulation of Guard Cell Turgor and Drought Tolerance by a Peroxisomal Acetate-Malate Shunt. Mol. Plant, 11, 1278–1291. Available at: 10.1016/j.molp.2018.07.008.

28. Dowle, Srinivasan, Gorecki and Chirico Package #x201C;data. table.” Extension of ‘data. Available at: ftp://ftp.musicbrainz.org/pub/cran/web/packages/data.table/data.table.pdf.

29. Fischer, D.S., Theis, F.J. and Yosef, N. (2018) Impulse model-based differential expression analysis of time course sequencing data. Nucleic Acids Res., 46, e119. Available at: 10.1093/nar/gky675.

30. Gallagher, J.P., Grover, C.E., Hu, G., Jareczek, J.J. and Wendel, J.F. (2020) Conservation and Divergence in Duplicated Fiber Coexpression Networks Accompanying Domestication of the Polyploid Gossypium hirsutum L. G3, 10, 2879–2892. Available at: 10.1534/g3.120.401362.

31. Gamblin, LeGendre, Collette, Lee, Moody, de Supinski and Futral (2015) The Spack package manager: bringing order to HPC software chaos. In SC15: International Conference for High-Performance Computing, Networking, Storage and Analysis. pp. 1–12. Available at: 10.1145/2807591.2807623.

32. Gilbert, M.K., Kim, H.J., Tang, Y., Naoumkina, M. and Fang, D.D. (2014) Comparative transcriptome analysis of short fiber mutants Ligon-lintless 1 and 2 reveals common mechanisms pertinent to fiber elongation in cotton (Gossypium hirsutum L.). PLoS One, 9, e95554. Available at: 10.1371/journal.pone.0095554.

33. Goodstein, D.M., Shu, S., Howson, R., et al. (2012) Phytozome: a comparative platform for green plant genomics. Nucleic Acids Res., 40, D1178–86. Available at: 10.1093/nar/gkr944.

34. Greenfield, A., Madar, A., Ostrer, H. and Bonneau, R. (2010) DREAM4: Combining genetic and dynamic information to identify biological networks and dynamical models. PLoS One, 5, e13397. Available at: 10.1371/journal.pone.0013397.

35. Haigler, C.H., Betancur, L., Stiff, M.R. and Tuttle, J.R. (2012) Cotton fiber: a powerful single-cell model for cell wall and cellulose research. Front. Plant Sci., 3, 104. Available at: 10.3389/fpls.2012.00104.

36. Haigler, C.H. and Roberts, A.W. (2019) Structure/function relationships in the rosette cellulose synthesis complex illuminated by an evolutionary perspective. Cellulose, 26, 227–247. Available at: 10.1007/s10570-018-2157-9.

37. Hovav, R., Chaudhary, B., Udall, J.A., Flagel, L. and Wendel, J.F. (2008) Parallel domestication, convergent evolution and duplicated gene recruitment in allopolyploid cotton. Genetics, 179, 1725–1733. Available at: 10.1534/genetics.108.089656.

38. Hovav, R., Udall, J.A., Hovav, E., Rapp, R., Flagel, L. and Wendel, J.F. (2008) A majority of cotton genes are expressed in single-celled fiber. Planta, 227, 319–329. Available at: 10.1007/s00425-007-0619-7.

39. Huang, G., Huang, J.-Q., Chen, X.-Y. and Zhu, Y.-X. (2021) Recent Advances and Future Perspectives in Cotton Research. Annu. Rev. Plant Biol., 72, 437–462. Available at: 10.1146/annurev-arplant-080720-113241.

40. Hu, G., Grover, C.E., Arick, M.A., Liu, M., Peterson, D.G. and Wendel, J.F. (2021) Homoeologous gene expression and co-expression network analyses and evolutionary inference in allopolyploids. Brief. Bioinform., 22, 1819–1835. Available at: 10.1093/bib/bbaa035.

41. Hu, G., Grover, C.E., Yuan, D., Dong, Y., Miller, E., Conover, J.L. and Wendel, J.F. (2021) Evolution and Diversity of the Cotton Genome. In M.-U.- Rahman, Y. Zafar, and T. Zhang, eds. Cotton Precision Breeding. Cham: Springer International Publishing, pp. 25–78. Available at: 10.1007/978-3-030-64504-5_2.

42. Hu, H., He, X., Tu, L., Zhu, L., Zhu, S., Ge, Z. and Zhang, X. (2016) GhJAZ2 negatively regulates cotton fiber initiation by interacting with the R2R3-MYB transcription factor GhMYB25-like. Plant J., 88, 921–935. Available at: https://onlinelibrary.wiley.com/doi/10.1111/tpj.13273.

43. Huynh-Thu, V.A., Irrthum, A., Wehenkel, L. and Geurts, P. (2010) Inferring regulatory networks from expression data using tree-based methods. PLoS One, 5. Available at: 10.1371/journal.pone.0012776.

44. Islam, M.S., Fang, D.D., Thyssen, G.N., Delhom, C.D., Liu, Y. and Kim, H.J. (2016) Comparative fiber property and transcriptome analyses reveal key genes potentially related to high fiber strength in cotton (Gossypium hirsutum L.) line MD52ne. BMC Plant Biol., 16, 36. Available at: 10.1186/s12870-016-0727-2.

45. Jareczek, J.J., Grover, C.E., Hu, G., Xiong, X., Arick, M.A., Ii, Peterson, D.G. and Wendel, J.F. (2023) Domestication over Speciation in Allopolyploid Cotton Species: A Stronger Transcriptomic Pull. Genes, 14. Available at: 10.3390/genes14061301.

46. Jareczek, J.J., Grover, C.E. and Wendel, J.F. (2023) Cotton fiber as a model for understanding shifts in cell development under domestication. Front. Plant Sci., 14, 1146802. Available at: 10.3389/fpls.2023.1146802.

47. Jensen, K.H., Berg-Sørensen, K., Bruus, H., Holbrook, N.M., Liesche, J., Schulz, A., Zwieniecki, M.A. and Bohr, T. (2016) Sap flow and sugar transport in plants. Rev. Mod. Phys., 88, 035007. Available at: https://link.aps.org/doi/10.1103/RevModPhys.88.035007.

48. Jiang, X., Fan, L., Li, P., Zou, X., Zhang, Z., Fan, S., Gong, J., Yuan, Y. and Shang, H. (2021) Co-expression network and comparative transcriptome analysis for fiber initiation and elongation reveal genetic differences in two lines from upland cotton CCRI70 RIL population. PeerJ, 9, e11812. Available at: 10.7717/peerj.11812.

49. Jiao, Y., Long, Y., Xu, K., et al. (2023) Weighted Gene Co-Expression Network Analysis Reveals Hub Genes for Fuzz Development in Gossypium hirsutum. Genes, 14. Available at: 10.3390/genes14010208.

50. Jin, J., Tian, F., Yang, D.-C., Meng, Y.-Q., Kong, L., Luo, J. and Gao, G. (2017) PlantTFDB 4.0: toward a central hub for transcription factors and regulatory interactions in plants. Nucleic Acids Res., 45, D1040–D1045. Available at: 10.1093/nar/gkw982.

51. Jin, J., Zhang, H., Kong, L., Gao, G. and Luo, J. (2014) PlantTFDB 3.0: a portal for the functional and evolutionary study of plant transcription factors. Nucleic Acids Res., 42, D1182–7. Available at: 10.1093/nar/gkt1016.

52. Kay, S., Hahn, S., Marois, E., Hause, G. and Bonas, U. (2007) A bacterial effector acts as a plant transcription factor and induces a cell size regulator. Science, 318, 648–651. Available at: 10.1126/science.1144956.

53. Khachiyan, L.G. (1996) Rounding of Polytopes in the Real Number Model of Computation. Mathematics of OR, 21, 307–320. Available at: 10.1287/moor.21.2.307.

54. Kim, H.J. (2018) Cotton Fiber Biosynthesis. In D. D. Fang, ed. Cotton Fiber: Physics, Chemistry and Biology. Cham: Springer International Publishing, pp. 133–150. Available at: 10.1007/978-3-030-00871-0_7.

55. Kim, H.J. (2015) Fiber Biology. In Agronomy Monographs. Madison, WI, USA: American Society of Agronomy, Inc., Crop Science Society of America, Inc., and Soil Science Society of America, Inc., pp. 97–127. Available at: http://doi.wiley.com/10.2134/agronmonogr57.2013.0022.

56. Kim, H.J., Thyssen, G.N., Song, X., Delhom, C.D. and Liu, Y. (2019) Functional divergence of cellulose synthase orthologs in between wild Gossypium raimondii and domesticated G. arboreum diploid cotton species. Cellulose. Available at: 10.1007/s10570-019-02744-y.

57. Kim, H.J. and Triplett, B.A. (2001) Cotton fiber growth in planta and in vitro. Models for plant cell elongation and cell wall biogenesis. Plant Physiol., 127, 1361–1366. Available at: https://www.ncbi.nlm.nih.gov/pubmed/11743074.

58. Kim, W.-C., Ko, J.-H., Kim, J.-Y., Kim, J., Bae, H.-J. and Han, K.-H. (2013) MYB46 directly regulates the gene expression of secondary wall-associated cellulose synthases in Arabidopsis. Plant J., 73, 26–36. Available at: 10.1111/j.1365-313x.2012.05124.x.

59. Kohel, R.J., Richmond, T.R. and Lewis, C.F. (1970) Texas Marker-1. Description of a Genetic Standard for Gossypium hirsutum L.1. Crop Sci., 10, 670–671. Available at: https://acsess.onlinelibrary.wiley.com/doi/10.2135/cropsci1970.0011183X001000060019x.

60. Kopka, J., Provart, N.J. and Müller-Röber, B. (1997) Potato guard cells respond to drying soil by a complex change in the expression of genes related to carbon metabolism and turgor regulation. Plant J., 11, 871–882. Available at: 10.1046/j.1365-313x.1997.11040871.x.

61. Kuanyshev, N., Deewan, A., Jagtap, S.S., Liu, J., Selvam, B., Chen, L.-Q., Shukla, D., Rao, C.V. and Jin, Y.-S. (2021) Identification and analysis of sugar transporters capable of co-transporting glucose and xylose simultaneously. Biotechnol. J., 16, e2100238. Available at: 10.1002/biot.202100238.

62. Langfelder, P. and Horvath, S. (2008) WGCNA: an R package for weighted correlation network analysis. BMC Bioinformatics, 9, 559. Available at: 10.1186/1471-2105-9-559.

63. Lee, C.M., Kafle, K., Belias, D.W., Park, Y.B., Glick, R.E., Haigler, C.H. and Kim, S.H. (2015) Comprehensive analysis of cellulose content, crystallinity, and lateral packing in Gossypium hirsutum and Gossypium barbadense cotton fibers using sum frequency generation, infrared and Raman spectroscopy, and X-ray diffraction. Cellulose, 22, 971–989. Available at: 10.1007/s10570-014-0535-5.

64. Lee, Y. and Szymanski, D.B. (2021) Multimerization variants as potential drivers of neofunctionalization. Sci Adv, 7. Available at: 10.1126/sciadv.abf0984.

65. Li, D.-D., Ruan, X.-M., Zhang, J., Wu, Y.-J., Wang, X.-L. and Li, X.-B. (2013) Cotton plasma membrane intrinsic protein 2s (PIP2s) selectively interact to regulate their water channel activities and are required for fibre development. New Phytol., 199, 695–707. Available at: 10.1111/nph.12309.

66. Li, J., Ruan, Y.-L., Dai, F., Zhu, S. and Zhang, T. (2023) Co-expression networks regulating cotton fiber initiation generated by comparative transcriptome analysis between fiberless XZ142FLM and GhVIN1i. Ind. Crops Prod., 194, 116323. Available at: https://www.sciencedirect.com/science/article/pii/S0926669023000870.

67. Li, S., Bashline, L., Lei, L. and Gu, Y. (2014) Cellulose synthesis and its regulation. Arabidopsis Book, 12, e0169. Available at: 10.1199/tab.0169.

68. Li, S.F., Milliken, O.N., Pham, H., Seyit, R., Napoli, R., Preston, J., Koltunow, A.M. and Parish, R.W. (2009) The Arabidopsis MYB5 transcription factor regulates mucilage synthesis, seed coat development, and trichome morphogenesis. Plant Cell, 21, 72–89. Available at: 10.1105/tpc.108.063503.

69. Liu, X., Qin, T., Ma, Q., Sun, J., Liu, Z., Yuan, M. and Mao, T. (2013) Light-regulated hypocotyl elongation involves proteasome-dependent degradation of the microtubule regulatory protein WDL3 in Arabidopsis. Plant Cell, 25, 1740–1755. Available at: 10.1105/tpc.113.112789.

70. Liu, Y., Ji, X., Nie, X., et al. (2015) Arabidopsis AtbHLH112 regulates the expression of genes involved in abiotic stress tolerance by binding to their E-box and GCG-box motifs. New Phytol., 207, 692–709. Available at: 10.1111/nph.13387.

71. Liu, Z., Sun, Z., Ke, H., et al. (2023) Transcriptome, Ectopic Expression and Genetic Population Analysis Identify Candidate Genes for Fiber Quality Improvement in Cotton. Int. J. Mol. Sci., 24. Available at: 10.3390/ijms24098293.

72. Li, X.-B., Fan, X.-P., Wang, X.-L., Cai, L. and Yang, W.-C. (2005) The cotton ACTIN1 gene is functionally expressed in fibers and participates in fiber elongation. Plant Cell, 17, 859–875. Available at: 10.1105/tpc.104.029629.

73. Li, Z., Wang, P., You, C., et al. (2020) Combined GWAS and eQTL analysis uncovers a genetic regulatory network orchestrating the initiation of secondary cell wall development in cotton. New Phytol., 226, 1738–1752. Available at: 10.1111/nph.16468.

74. Lockhart, J.A. (1965) An analysis of irreversible plant cell elongation. J. Theor. Biol., 8, 264–275. Available at: 10.1016/0022-5193(65)90077-9.

75. Love, M.I., Huber, W. and Anders, S. (2014) Moderated estimation of fold change and dispersion for RNA-seq data with DESeq2. Genome Biol., 15, 550. Available at: 10.1186/s13059-014-0550-8.

76. Luo, M., Xiao, Y., Li, X., et al. (2007) GhDET2, a steroid 5alpha-reductase, plays an important role in cotton fiber cell initiation and elongation. Plant J., 51, 419–430. Available at: https://onlinelibrary.wiley.com/doi/10.1111/j.1365-313X.2007.03144.x.

77. Machado, A., Wu, Y., Yang, Y., Llewellyn, D.J. and Dennis, E.S. (2009) The MYB transcription factor GhMYB25 regulates early fibre and trichome development. Plant J., 59, 52–62. Available at: 10.1111/j.1365-313X.2009.03847.x.

78. MacMillan, C.P., Birke, H., Chuah, A., Brill, E., Tsuji, Y., Ralph, J., Dennis, E.S., Llewellyn, D. and Pettolino, F.A. (2017) Tissue and cell-specific transcriptomes in cotton reveal the subtleties of gene regulation underlying the diversity of plant secondary cell walls. BMC Genomics, 18, 539. Available at: 10.1186/s12864-017-3902-4.

79. MacRobbie, E.A.C. (2006) Control of volume and turgor in stomatal guard cells. J. Membr. Biol., 210, 131–142. Available at: 10.1007/s00232-005-0851-7.

80. Madeira, F., Pearce, M., Tivey, A.R.N., Basutkar, P., Lee, J., Edbali, O., Madhusoodanan, N., Kolesnikov, A. and Lopez, R. (2022) Search and sequence analysis tools services from EMBL-EBI in 2022. Nucleic Acids Res., 50, W276–W279. Available at: 10.1093/nar/gkac240.

81. Mao, G., Buschmann, H., Doonan, J.H. and Lloyd, C.W. (2006) The role of MAP65-1 in microtubule bundling during Zinnia tracheary element formation. J. Cell Sci., 119, 753–758. Available at: 10.1242/jcs.02813.

82. Ma, Q., Wang, N., Hao, P., et al. (2019) Genome-wide identification and characterization of TALE superfamily genes in cotton reveals their functions in regulating secondary cell wall biosynthesis. BMC Plant Biol., 19, 432. Available at: 10.1186/s12870-019-2026-1.

83. Marbach, D., Costello, J.C., Küffner, R., et al. (2012) Wisdom of crowds for robust gene network inference. Nat. Methods, 9, 796–804. Available at: 10.1038/nmeth.2016.

84. McBride, Z., Chen, D., Reick, C., Xie, J. and Szymanski, D.B. (2017) Global Analysis of Membrane-associated Protein Oligomerization Using Protein Correlation Profiling. Mol. Cell. Proteomics, 16, 1972–1989. Available at: 10.1074/mcp.RA117.000276.

85. McFarlane, H.E., Mutwil-Anderwald, D., Verbančič, J., et al. (2021) A G protein-coupled receptor-like module regulates cellulose synthase secretion from the endomembrane system in Arabidopsis. Dev. Cell, 56, 1484–1497.e7. Available at: 10.1016/j.devcel.2021.03.031.

86. Meinert, M.C. and Delmer, D.P. (1977) Changes in biochemical composition of the cell wall of the cotton fiber during development. Plant Physiol., 59, 1088–1097. Available at: 10.1104/pp.59.6.1088.

87. Naoumkina, M., Thyssen, G.N. and Fang, D.D. (2015) RNA-seq analysis of short fiber mutants Ligon-lintless-1 (Li 1) and -2 (Li 2) revealed important role of aquaporins in cotton (Gossypium hirsutum L.) fiber elongation. BMC Plant Biol., 15, 65. Available at: 10.1186/s12870-015-0454-0.

88. Page, J.T., Gingle, A.R. and Udall, J.A. (2013) PolyCat: a resource for genome categorization of sequencing reads from allopolyploid organisms. G3, 3, 517–525. Available at: 10.1534/g3.112.005298.

89. Paterson, A.H., Wendel, J.F., Gundlach, H., et al. (2012) Repeated polyploidization of Gossypium genomes and the evolution of spinnable cotton fibres. Nature, 492, 423–427. Available at: 10.1038/nature11798.

90. Pedersen, G.B., Blaschek, L., Frandsen, K.E.H., Noack, L.C. and Persson, S. (2023) Cellulose synthesis in land plants. Mol. Plant, 16, 206–231. Available at: 10.1016/j.molp.2022.12.015.

91. Pedersen, T.L. (2022) ggforce: Accelerating “ggplot2,” Available at: https://ggforce.data-imaginist.com, https://github.com/thomasp85/ggforce.

92. Perez-Riverol, Y., Bai, J., Bandla, C., et al. (2022) The PRIDE database resources in 2022: a hub for mass spectrometry-based proteomics evidences. Nucleic Acids Res., 50, D543–D552. Available at: 10.1093/nar/gkab1038.

93. Pettolino, F.A., Yulia, D., Bacic, A. and Llewellyn, D.J. (2022) Polysaccharide composition during cotton seed fibre development: temporal differences between species and in different seasons. Journal of Cotton Research, 5, 1–13. Available at: https://jcottonres.biomedcentral.com/articles/10.1186/s42397-022-00136-5 [Accessed December 21, 2022].

94. Potikha, T.S., Collins, C.C., Johnson, D.I., Delmer, D.P. and Levine, A. (1999) The involvement of hydrogen peroxide in the differentiation of secondary walls in cotton fibers. Plant Physiol., 119, 849–858. Available at: 10.1104/pp.119.3.849.

95. Proseus, T.E., Zhu, G.L. and Boyer, J.S. (2000) Turgor, temperature and the growth of plant cells: using Chara corallina as a model system. J. Exp. Bot., 51, 1481–1494. Available at: 10.1093/jexbot/51.350.1481.

96. Pu, L., Li, Q., Fan, X., Yang, W. and Xue, Y. (2008) The R2R3 MYB Transcription Factor GhMYB109 Is Required for Cotton Fiber Development. Genetics, 180, 811–820. Available at: https://academic.oup.com/genetics/article-abstract/180/2/811/6073815 [Accessed April 12, 2022].

97. Qin, Y.-M. and Zhu, Y.-X. (2011) How cotton fibers elongate: a tale of linear cell-growth mode. Curr. Opin. Plant Biol., 14, 106–111. Available at: 10.1016/j.pbi.2010.09.010.

99. Qin, Y., Sun, H., Hao, P., Wang, H., Wang, C., Ma, L., Wei, H. and Yu, S. (2019) Transcriptome analysis reveals differences in the mechanisms of fiber initiation and elongation between long- and short-fiber cotton (Gossypium hirsutum L.) lines. BMC Genomics, 20, 633. Available at: 10.1186/s12864-019-5986-5.

100. Qu, J., Ye, J., Geng, Y.-F., Sun, Y.-W., Gao, S.-Q., Zhang, B.-P., Chen, W. and Chua, N.-H. (2012) Dissecting functions of KATANIN and WRINKLED1 in cotton fiber development by virus-induced gene silencing. Plant Physiol., 160, 738–748. Available at: 10.1104/pp.112.198564.

101. Rapp, R.A., Haigler, C.H., Flagel, L., Hovav, R.H., Udall, J.A. and Wendel, J.F. (2010) Gene expression in developing fibres of Upland cotton (Gossypium hirsutum L.) was massively altered by domestication. BMC Biol., 8, 139. Available at: 10.1186/1741-7007-8-139.

102. R Core Team (2022) R: A language and environment for statistical computing, Vienna, Austria: R Foundation for Statistical Computing. Available at: https://www.R-project.org/.

103. Rhodes, D. and Samaras, Y. (2020) Genetic control of osmoregulation in plants. Am. J. Physiol. Lung Cell. Mol. Physiol., 347–361. Available at: https://www.taylorfrancis.com/chapters/edit/10.1201/9780367812140-25/genetic-control-osmoregulation-plants-david-rhodes-yiannis-samaras.

104. Richmond, T.A. and Somerville, C.R. (2000) The cellulose synthase superfamily. Plant Physiol., 124, 495–498. Available at: 10.1104/pp.124.2.495.

105. Rosvall, M. and Bergstrom, C.T. (2008) Maps of random walks on complex networks reveal community structure. Proc. Natl. Acad. Sci. U. S. A., 105, 1118–1123. Available at: 10.1073/pnas.0706851105.

106. Ruan, Y.-L. (2005) Recent advances in understanding cotton fibre and seed development. Seed Sci. Res., 15, 269–280. Available at: 10.1079/SSR2005217 [Accessed March 5, 2024].

107. Ruan, Y.L., Llewellyn, D.J. and Furbank, R.T. (2001) The control of single-celled cotton fiber elongation by developmentally reversible gating of plasmodesmata and coordinated expression of sucrose and K+ transporters and expansin. Plant Cell, 13, 47–60. Available at: 10.1105/tpc.13.1.47.

108. Ruan, Y.L., Mate, C., Patrick, J.W. and Brady, C.J. (1995) Non-Destructive Collection of Apoplast Fluid From Developing Tomato Fruit Using a Pressure Dehydration Procedure. Funct. Plant Biol., 22, 761–769. Available at: 10.1071/pp9950761 [Accessed April 24, 2024].

109. Ryser, U. (1977) Cell wall growth in elongating cotton fibers: an autoradiographic study. Cytobiologie.

110. Schiffthaler, B., Zalen, E. van Serrano, A.R., Street, N.R. and Delhomme, N. (2023) Seiðr: Efficient calculation of robust ensemble gene networks. Heliyon, 9, e16811. Available at: https://www.sciencedirect.com/science/article/pii/S2405844023040185.

111. Schneider, R., Klooster, K.V., Picard, K.L., et al. (2021) Long-term single-cell imaging and simulations of microtubules reveal principles behind wall patterning during proto-xylem development. Nat. Commun., 12, 669. Available at: 10.1038/s41467-021-20894-1.

112. Schubert, A.M., Benedict, C.R., Berlin, J.D. and Kohel, R.J. (1973) Cotton fiber development-kinetics of cell elongation and secondary wall thickening 1. Crop Sci., 13, 704–709. Available at: 10.2135/cropsci1973.0011183x001300060035x.

113. Seagull, R.W. (1993) Cytoskeletal involvement in cotton fiber growth and development. Micron, 24, 643–660. Available at: https://www.sciencedirect.com/science/article/pii/096843289390042Y.

114. Shan, C.-M., Shangguan, X.-X., Zhao, B., et al. (2014) Control of cotton fibre elongation by a homeodomain transcription factor GhHOX3. Nat. Commun., 5, 5519. Available at: 10.1038/ncomms6519.

115. Shannon, P., Markiel, A., Ozier, O., Baliga, N.S., Wang, J.T., Ramage, D., Amin, N., Schwikowski, B. and Ideker, T. (2003) Cytoscape: a software environment for integrated models of biomolecular interaction networks. Genome Res., 13, 2498–2504. Available at: 10.1101/gr.1239303.

116. Singh, B., Avci, U., Eichler Inwood, S.E., et al. (2009) A specialized outer layer of the primary cell wall joins elongating cotton fibers into tissue-like bundles. Plant Physiol., 150, 684–699. Available at: https://academic.oup.com/plphys/article-abstract/150/2/684/6107960.

117. Smart, L.B., Vojdani, F., Maeshima, M. and Wilkins, T.A. (1998) Genes involved in osmoregulation during turgor-driven cell expansion of developing cotton fibers are differentially regulated. Plant Physiol., 116, 1539–1549. Available at: 10.1104/pp.116.4.1539.

118. Steudle, E. and Zimmermann, U. (1977) Effect of turgor pressure and cell size on the wall elasticity of plant cells. Plant Physiol., 59, 285–289. Available at: 10.1104/pp.59.2.285.

119. Stiff, M.R. and Haigler, C.H. (2016) Cotton fiber tips have diverse morphologies and show evidence of apical cell wall synthesis. Sci. Rep., 6, 27883. Available at: 10.1038/srep27883.

120. Sun, W., Gao, Z., Wang, J., et al. (2019) Cotton fiber elongation requires the transcription factor GhMYB212 to regulate sucrose transportation into expanding fibers. New Phytol., 222, 864–881. Available at: 10.1111/nph.15620.

121. Sun, X., Gong, S.-Y., Nie, X.-Y., Li, Y., Li, W., Huang, G.-Q. and Li, X.-B. (2015) A R2R3-MYB transcription factor that is specifically expressed in cotton (Gossypium hirsutum) fibers affects secondary cell wall biosynthesis and deposition in transgenic Arabidopsis. Physiol. Plant., 154, 420–432. Available at: https://onlinelibrary.wiley.com/doi/10.1111/ppl.12317.

122. Swaminathan, S., Grover, C.E., Mugisha, A.S., et al. (2024) Daily glycome and transcriptome profiling reveals polysaccharide structures and glycosyltransferases critical for cotton fiber growth. bioRxiv, 2024.04.23.589927. Available at: https://www.biorxiv.org/content/10.1101/2024.04.23.589927v1 [Accessed May 10, 2024].

123. Takatsuka, H., Higaki, T. and Umeda, M. (2018) Actin Reorganization Triggers Rapid Cell Elongation in Roots. Plant Physiol., 178, 1130–1141. Available at: 10.1104/pp.18.00557.

124. Tange, O. (2022) GNU Parallel 20220522 (’NATO’), Available at: https://zenodo.org/record/6570228.

125. Tang, W., Tu, L., Yang, X., Tan, J., Deng, F., Hao, J., Guo, K., Lindsey, K. and Zhang, X. (2014) The calcium sensor GhCaM7 promotes cotton fiber elongation by modulating reactive oxygen species (ROS) production. New Phytol., 202, 509–520. Available at: https://onlinelibrary.wiley.com/doi/10.1111/nph.12676.

126. Thaker, V.S., Rabadia, V.S. and Singh, Y.D. (1999) Physiological and biochemical changes associated with cotton fibre development. Acta Physiol. Plant, 21, 57–61. Available at: 10.1007/s11738-999-0027-7.

127. Tian, F., Yang, D.-C., Meng, Y.-Q., Jin, J. and Gao, G. (2020) PlantRegMap: charting functional regulatory maps in plants. Nucleic Acids Res., 48, D1104–D1113. Available at: 10.1093/nar/gkz1020.

128. Tiwari, S.C. and Wilkins, T.A. (1995) Cotton (Gossypium hirsutum) seed trichomes expand via diffuse growing mechanism. Can. J. Bot., 73, 746–757. Available at: https://cdnsciencepub.com/doi/abs/10.1139/b95-081.

129. Tuttle, J.R., Nah, G., Duke, M.V., Alexander, D.C., Guan, X., Song, Q., Chen, Z.J., Scheffler, B.E. and Haigler, C.H. (2015) Metabolomic and transcriptomic insights into how cotton fiber transitions to secondary wall synthesis, represses lignification, and prolongs elongation. BMC Genomics, 16, 477. Available at: 10.1186/s12864-015-1708-9.

130. Viot, C.R. and Wendel, J.F. (2023) Evolution of the Cotton Genus, Gossypium, and Its Domestication in the Americas. CRC Crit. Rev. Plant Sci., 1–33. Available at: https://www.tandfonline.com/doi/abs/10.1080/07352689.2022.2156061.

131. Wang, L., Kartika, D. and Ruan, Y.-L. (2021) Looking into “hair tonics” for cotton fiber initiation. New Phytol., 229, 1844–1851. Available at: 10.1111/nph.16898.

132. Wang, N.-N., Li, Y., Chen, Y.-H., Lu, R., Zhou, L., Wang, Y., Zheng, Y. and Li, X.-B. (2021) Phosphorylation of WRKY16 by MPK3-1 is essential for its transcriptional activity during fiber initiation and elongation in cotton (Gossypium hirsutum). Plant Cell, 33, 2736–2752. Available at: 10.1093/plcell/koab153.

133. Wang, Q.Q., Liu, F., Chen, X.S., Ma, X.J., Zeng, H.Q. and Yang, Z.M. (2010) Transcriptome profiling of early developing cotton fiber by deep-sequencing reveals significantly differential expression of genes in a fuzzless/lintless mutant. Genomics, 96, 369–376. Available at: 10.1016/j.ygeno.2010.08.009.

134. Wickham, H. (2016) ggplot2: Elegant Graphics for Data Analysis, Springer-Verlag New York.

135. Wickham, H., Averick, M., Bryan, J., et al. (2019) Welcome to the tidyverse. J. Open Source Softw., 4, 1686. Available at: https://joss.theoj.org/papers/10.21105/joss.01686.

136. Worden, N., Wilkop, T.E., Esteve, V.E., et al. (2015) CESA TRAFFICKING INHIBITOR inhibits cellulose deposition and interferes with the trafficking of cellulose synthase complexes and their associated proteins KORRIGAN1 and POM2/CELLULOSE SYNTHASE INTERACTIVE PROTEIN1. Plant Physiol., 167, 381–393. Available at: 10.1104/pp.114.249003.

137. Xiao, G., Zhao, P. and Zhang, Y. (2019) A Pivotal Role of Hormones in Regulating Cotton Fiber Development. Front. Plant Sci., 10, 87. Available at: 10.3389/fpls.2019.00087.

138. Xu, Q. and Liesche, J. (2021) Sugar export from Arabidopsis leaves: actors and regulatory strategies. J. Exp. Bot., 72, 5275–5284. Available at: 10.1093/jxb/erab241.

139. Yanagisawa, M., Desyatova, A.S., Belteton, S.A., Mallery, E.L., Turner, J.A. and Szymanski, D.B. (2015) Patterning mechanisms of cytoskeletal and cell wall systems during leaf trichome morphogenesis. Nature Plants, 1, 1–8. Available at: https://www.nature.com/articles/nplants201514 [Accessed March 29, 2024].

140. Yanagisawa, M., Keynia, S., Belteton, S., Turner, J.A. and Szymanski, D. (2022) A conserved cellular mechanism for cotton fibre diameter and length control. in silico Plants, 4. Available at: https://academic.oup.com/insilicoplants/article-pdf/4/1/diac004/45932206/diac004.pdf [Accessed January 19, 2023].

141. Yang, Z., Zhang, C., Yang, X., et al. (2014) PAG1, a cotton brassinosteroid catabolism gene, modulates fiber elongation. New Phytol., 203, 437–448. Available at: 10.1111/nph.12824.

142. Yoo, M.-J. and Wendel, J.F. (2014) Comparative evolutionary and developmental dynamics of the cotton (Gossypium hirsutum) fiber transcriptome. PLoS Genet., 10, e1004073. Available at: 10.1371/journal.pgen.1004073.

143. Yu, Y., Wu, S., Nowak, J., et al. (2019) Live-cell imaging of the cytoskeleton in elongating cotton fibres. Nat Plants, 5, 498–504. Available at: 10.1038/s41477-019-0418-8.

144. Zabotina, O.A., Avci, U., Cavalier, D., Pattathil, S., Chou, Y.-H., Eberhard, S., Danhof, L., Keegstra, K. and Hahn, M.G. (2012) Mutations in multiple XXT genes of Arabidopsis reveal the complexity of xyloglucan biosynthesis. Plant Physiol., 159, 1367–1384. Available at: 10.1104/pp.112.198119.

145. Zhang, J., Xie, M., Tuskan, G.A., Muchero, W. and Chen, J.-G. (2018) Recent Advances in the Transcriptional Regulation of Secondary Cell Wall Biosynthesis in the Woody Plants. Front. Plant Sci., 9, 1535. Available at: 10.3389/fpls.2018.01535.

146. Zhang, X., Cao, J., Huang, C., Zheng, Z., Liu, X., Shangguan, X., Wang, L., Zhang, Y. and Chen, Z. (2021) Characterization of cotton ARF factors and the role of GhARF2b in fiber development. BMC Genomics, 22, 202. Available at: 10.1186/s12864-021-07504-6.

147. Zhang, X., Hu, D.-P., Li, Y., Chen, Y., Abidallha, E.H.M.A., Dong, Z.-D., Chen, D.-H. and Zhang, L. (2017) Developmental and hormonal regulation of fiber quality in two natural-colored cotton cultivars. J. Integr. Agric., 16, 1720–1729. Available at: https://www.sciencedirect.com/science/article/pii/S2095311916615046.

148. Zhang, Y., Yu, J., Wang, X., Durachko, D.M., Zhang, S. and Cosgrove, D.J. (2021) Molecular insights into the complex mechanics of plant epidermal cell walls. Science, 372, 706–711. Available at: 10.1126/science.abf2824.

149. Zhao, T., Xu, X., Wang, M., Li, C., Li, C., Zhao, R., Zhu, S., He, Q. and Chen, J. (2019) Identification and profiling of upland cotton microRNAs at fiber initiation stage under exogenous IAA application. BMC Genomics, 20, 421. Available at: 10.1186/s12864-019-5760-8.

150. Zhong, R., Allen, J.D., Xiao, G. and Xie, Y. (2014) Ensemble-based network aggregation improves the accuracy of gene network reconstruction. PLoS One, 9, e106319. Available at: 10.1371/journal.pone.0106319.

151. Zhong, R., Cui, D. and Ye, Z.-H. (2019) Secondary cell wall biosynthesis. New Phytol., 221, 1703–1723. Available at: 10.1111/nph.15537.

152. Zhong, R., Demura, T. and Ye, Z.-H. (2006) SND1, a NAC domain transcription factor, is a key regulator of secondary wall synthesis in fibers of Arabidopsis. Plant Cell, 18, 3158–3170. Available at: 10.1105/tpc.106.047399.

153. Zhou, X., Hu, W., Li, B., Yang, Y., Zhang, Y., Thow, K., Fan, L. and Qu, Y. (2019) Proteomic profiling of cotton fiber developmental transition from cell elongation to secondary wall deposition. Acta Biochim. Biophys. Sin., 51, 1168–1177. Available at: 10.1093/abbs/gmz111.

154. Zhu, H., Han, X., Lv, J., Zhao, L., Xu, X., Zhang, T. and Guo, W. (2011) Structure, expression differentiation and evolution of duplicated fiber developmental genes in Gossypium barbadense and G. hirsutum. BMC Plant Biol., 11, 40. Available at: 10.1186/1471-2229-11-40.

155. Zimmermann, U. (1978) Physics of Turgor- and Osmoregulation. Annu. Rev. Plant Biol., 29, 121–148. Available at: 10.1146/annurev.pp.29.060178.001005 [Accessed March 22, 2024].

156. Zou, X., Ali, F., Jin, S., Li, F. and Wang, Z. (2022) RNA-Seq with a novel glabrous-ZM24fl reveals some key lncRNAs and the associated targets in fiber initiation of cotton. BMC Plant Biol., 22, 61. Available at: 10.1186/s12870-022-03444-9.

