## Supplementary figures and images for "A high-resolution model of gene expression during *Gossypium hirsutum* (cotton) fiber development"

### Supplementary Figure 1

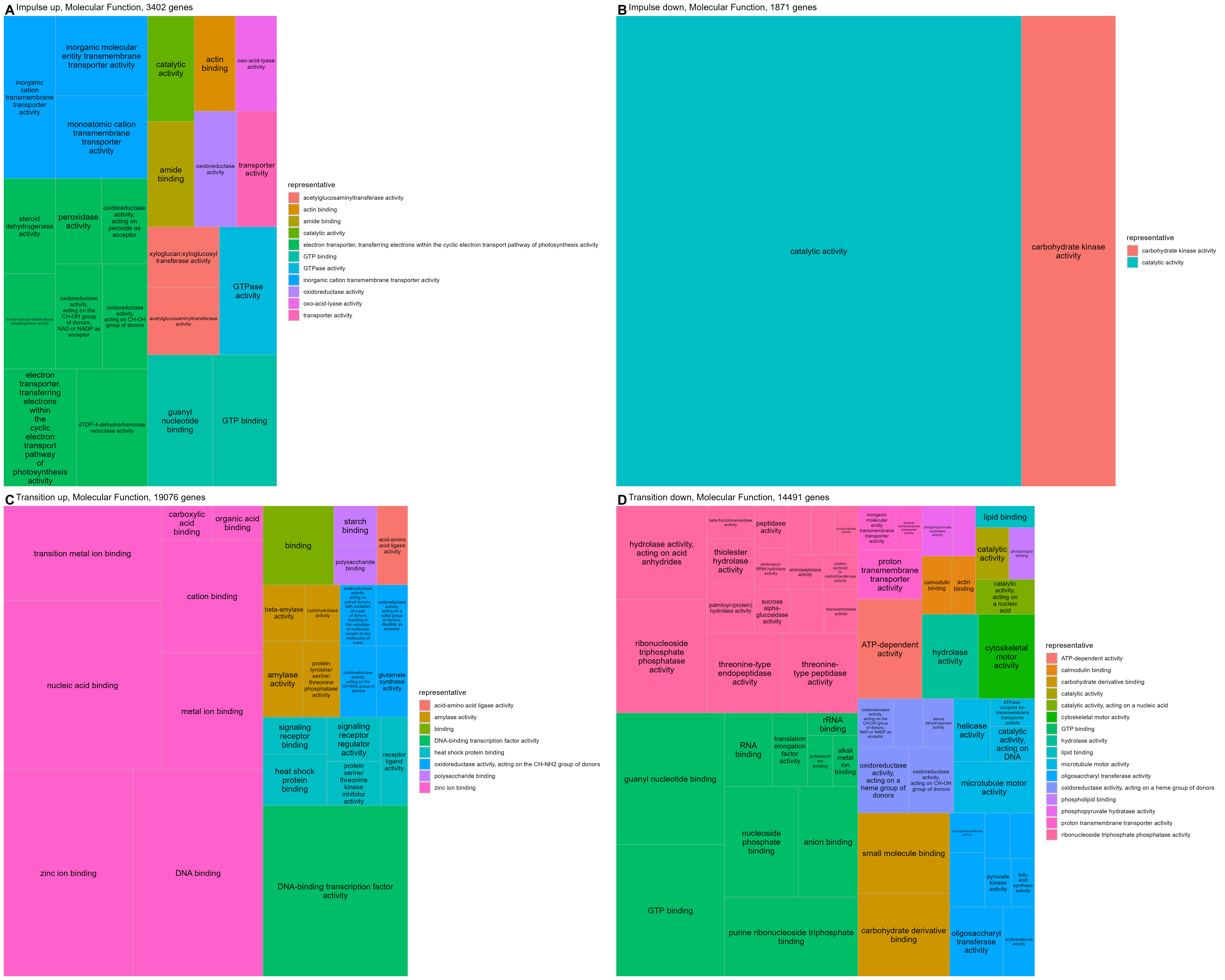

### Supplementary Figure 2

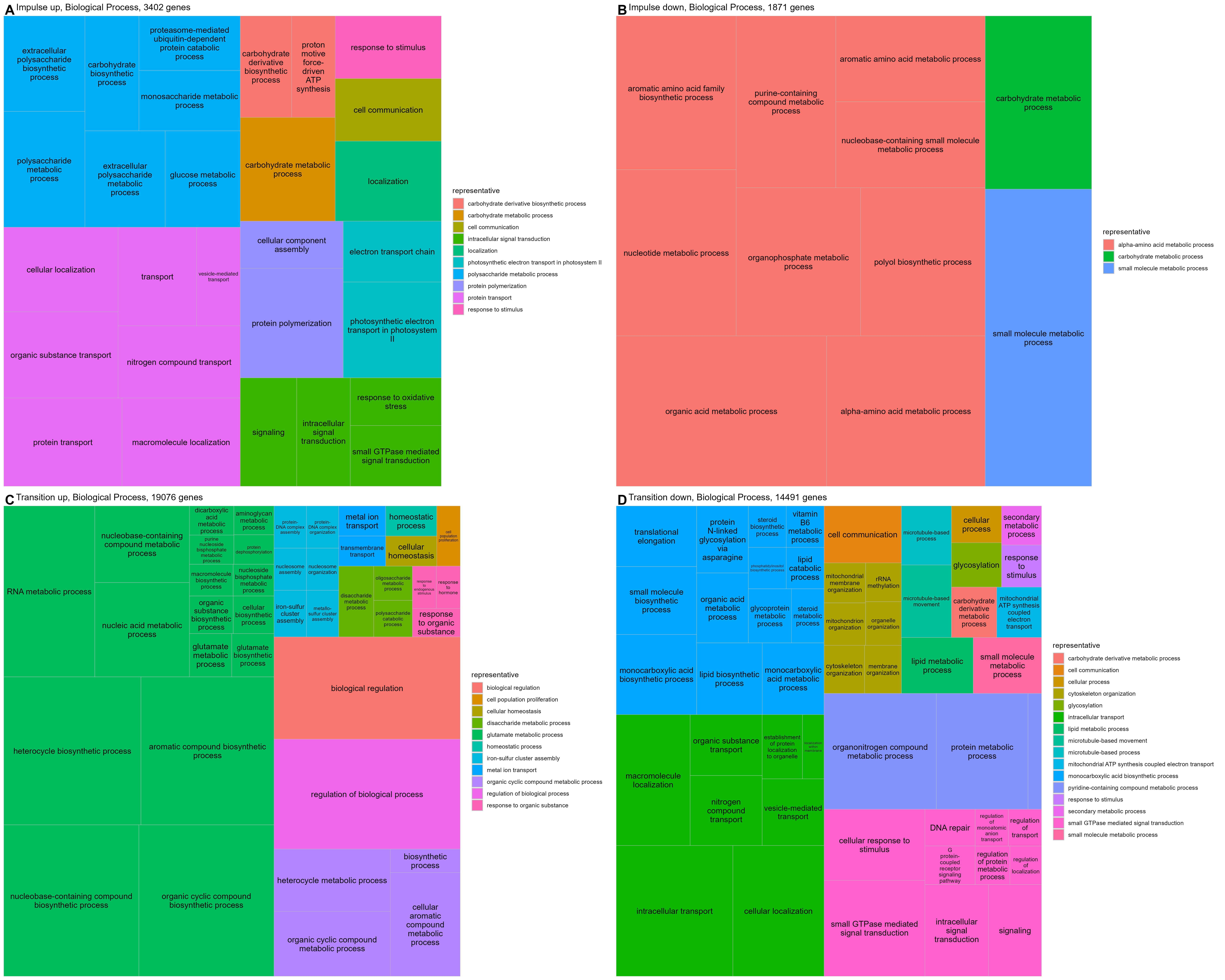

### Supplementary Figure 3

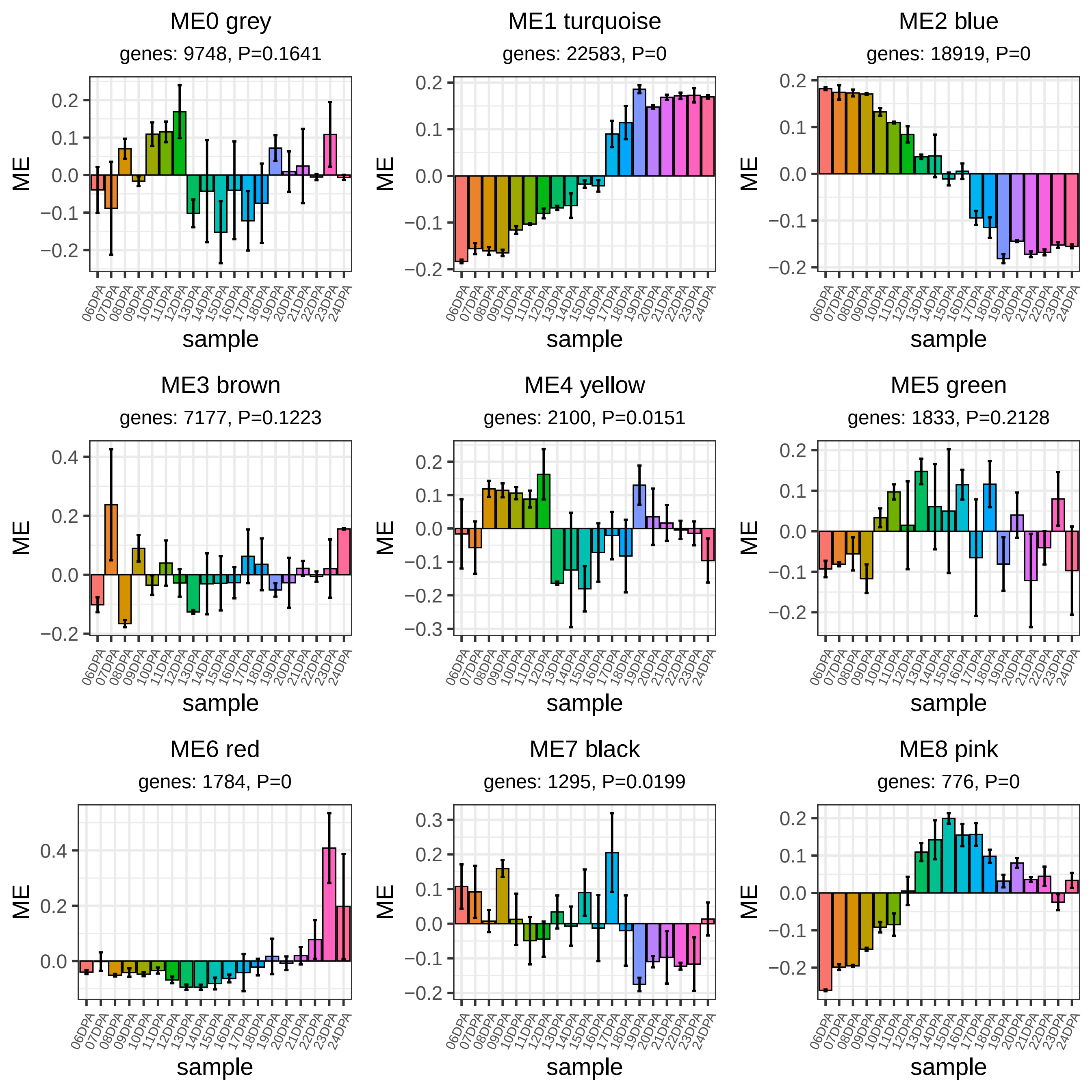

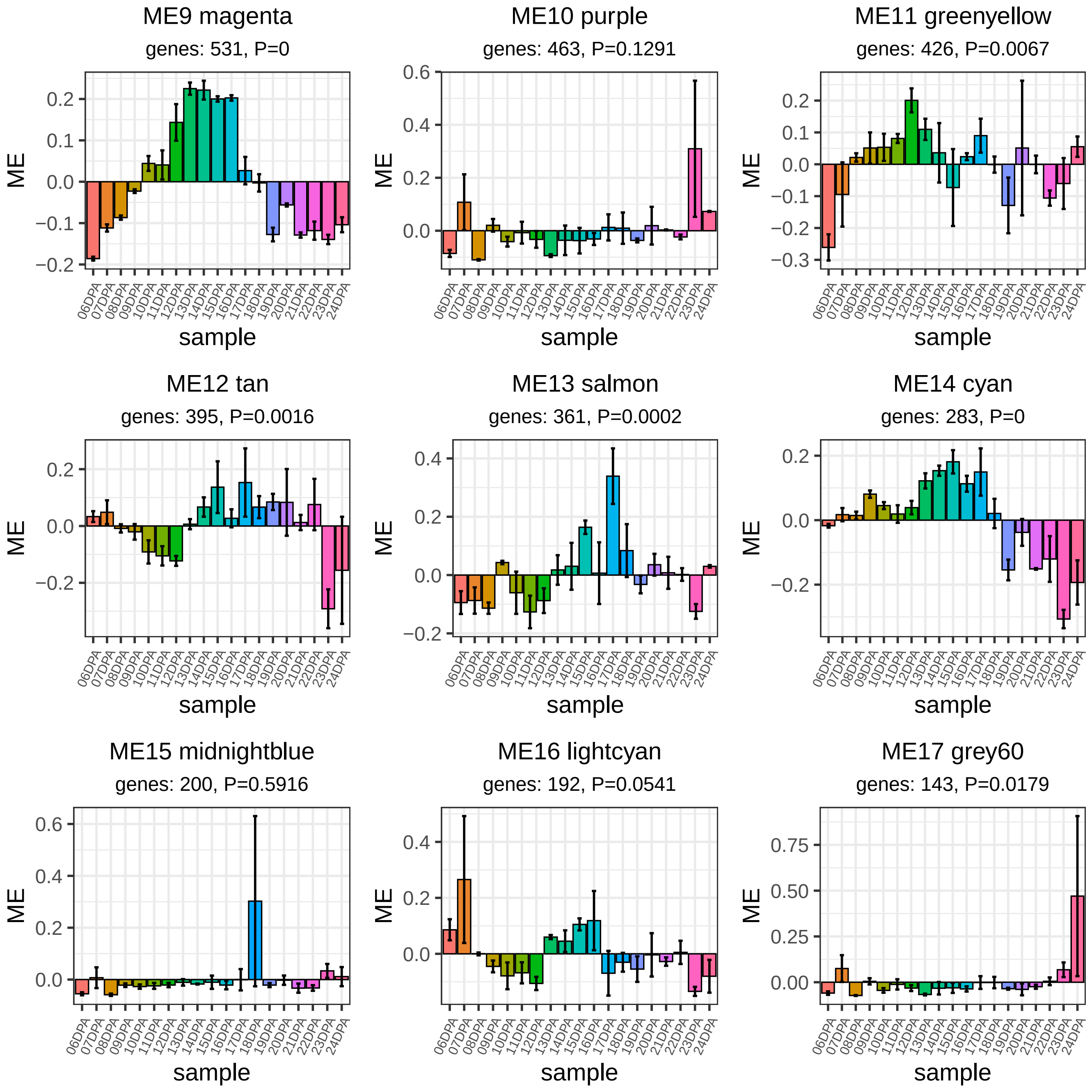

### Supplementary Figure 4

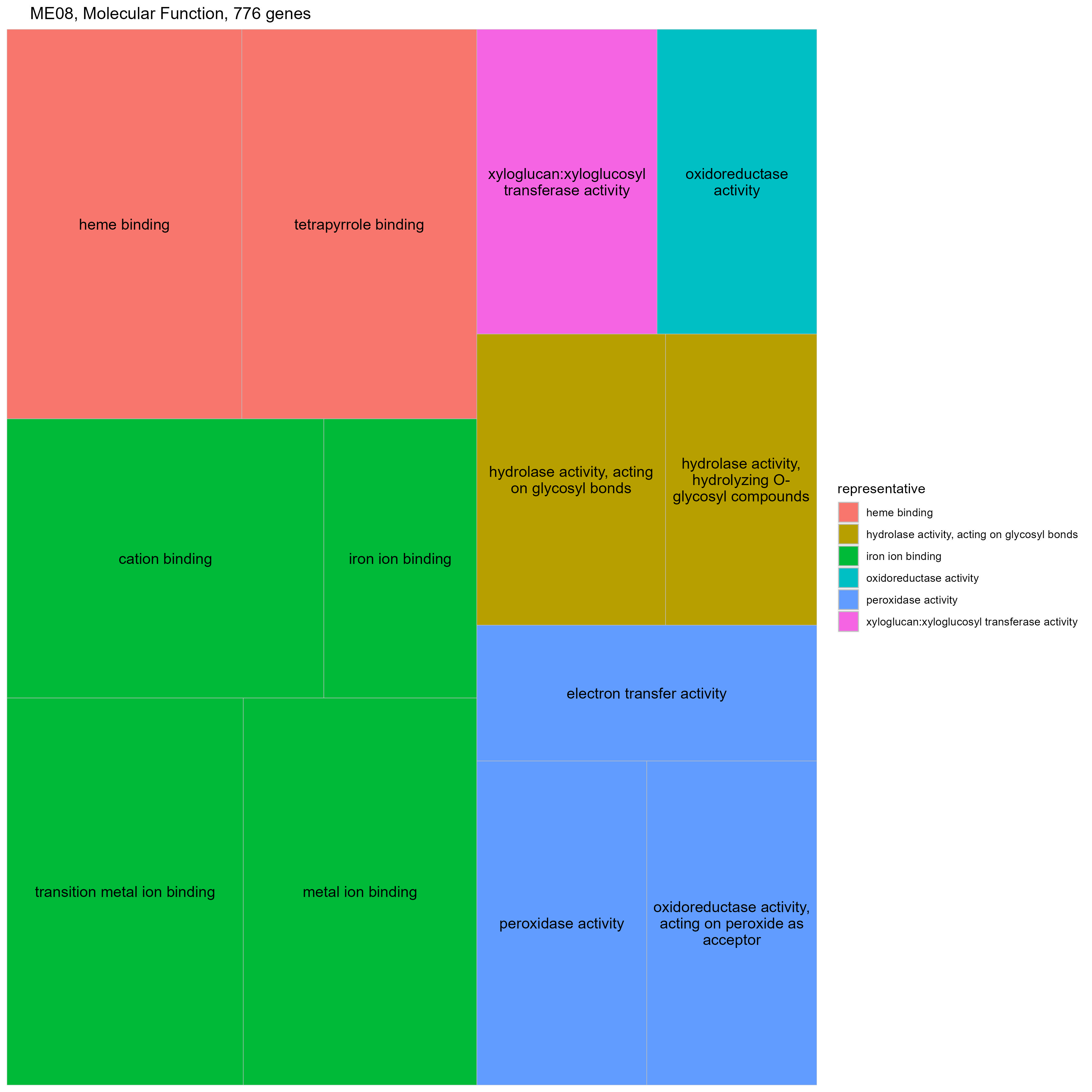

### Supplementary Figure 5

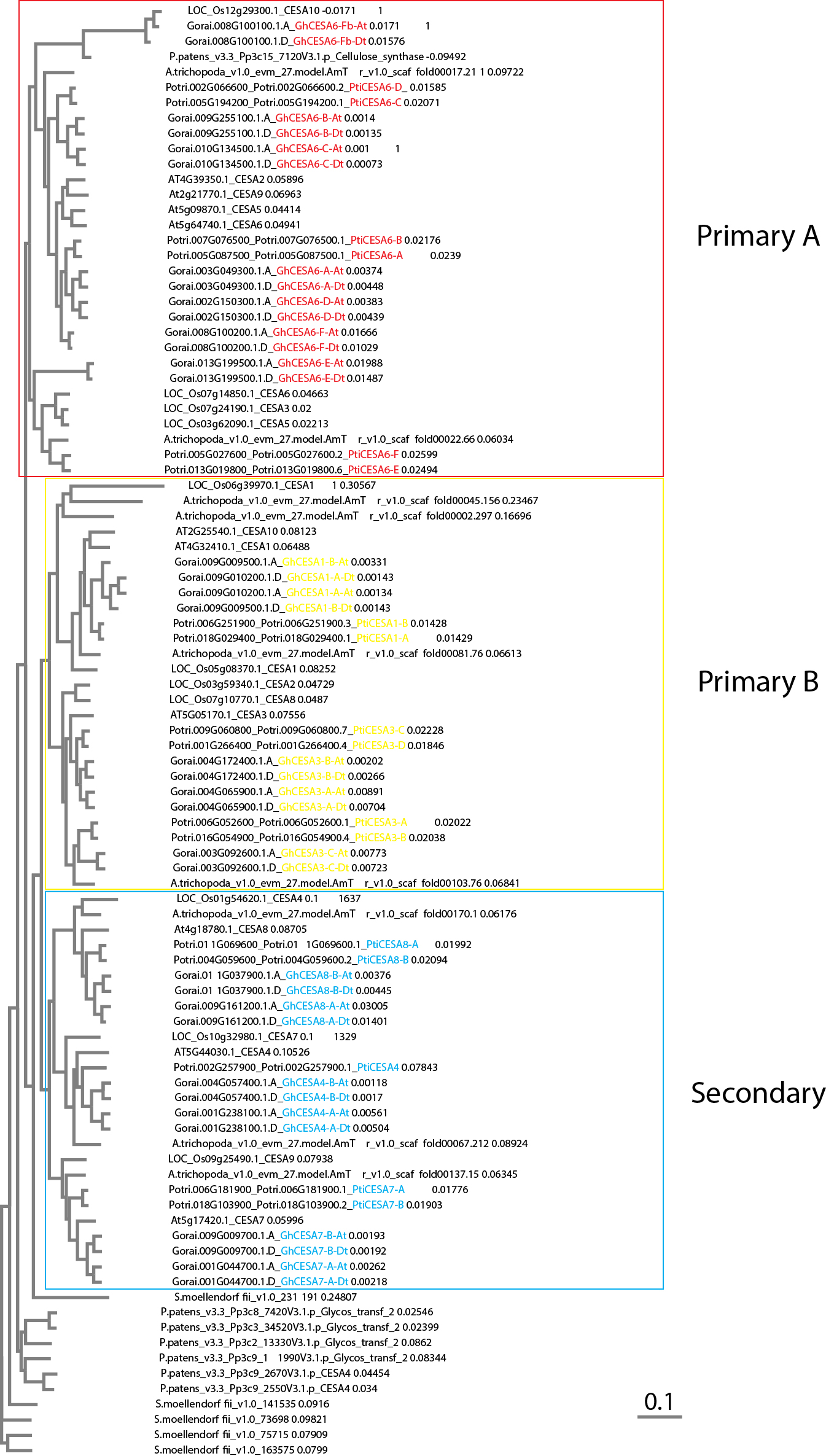

### Supplementary Figure 6

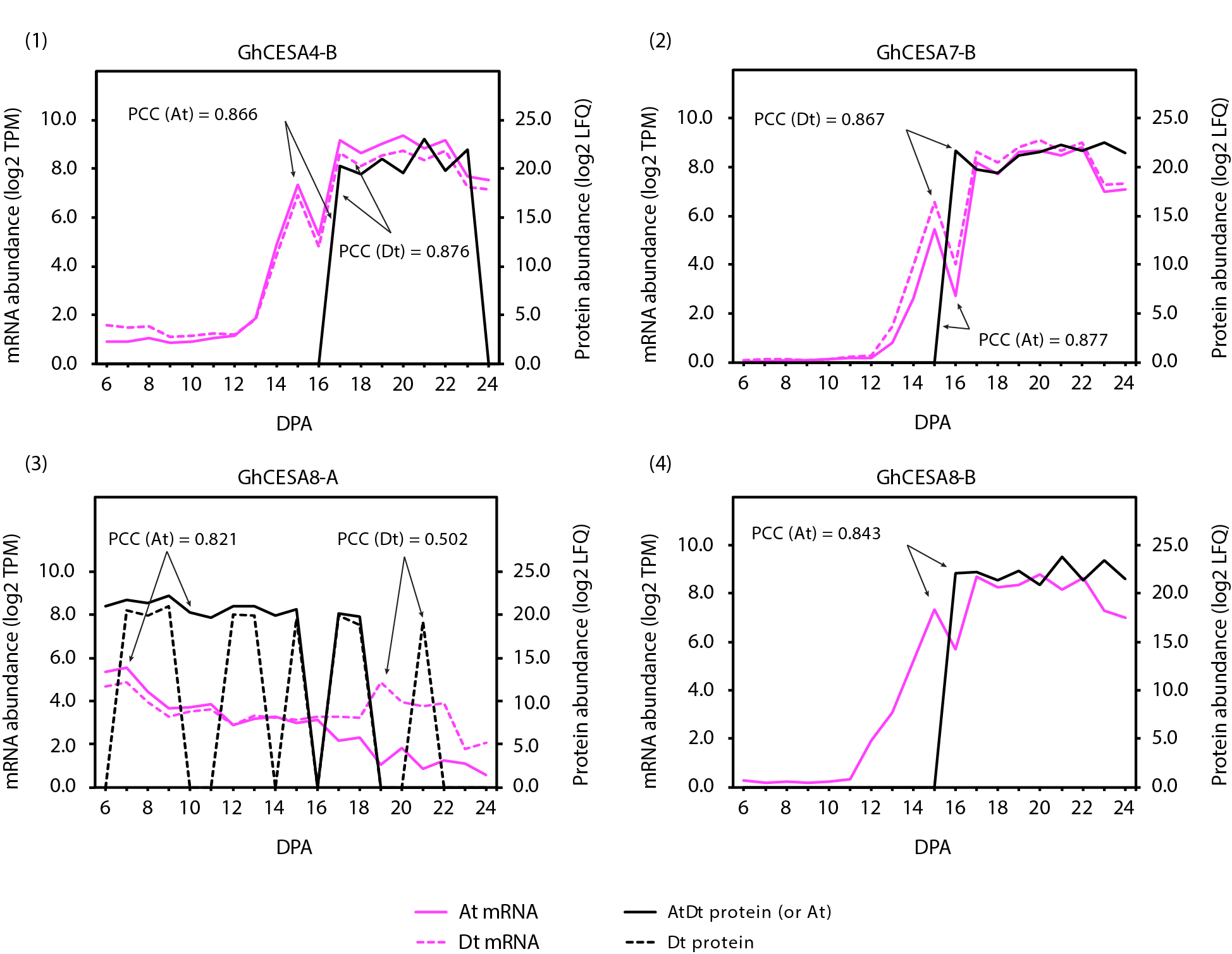
